# TRPM2 deficiency protects against atherosclerosis by inhibiting TRPM2-CD36 inflammatory axis in macrophages

**DOI:** 10.1101/2021.07.29.454234

**Authors:** Pengyu Zong, Jianlin Feng, Zhichao Yue, Albert S. Yu, Yasuo Mori, Lixia Yue

**Affiliations:** Department of Cell Biology, Calhoun Cardiology Center, University of Connecticut School of Medicine (UConn Health), Farmington, CT 06030, USA; Laboratory of Molecular Biology, Department of Synthetic Chemistry and Biological Chemistry, Graduate School of Engineering, Kyoto University, Katsura Campus A4-218 Nishikyo-ku, Kyoto 615-8510, Japan

**Author notes:** Corresponding author: Lixia Yue.

**Keywords:** Atherosclerosis, TRPM2, macrophages, CD36, oxLDL, TSP1

## Abstract

Atherosclerosis is the major cause of ischemic heart diseases and ischemic brain stroke, which are the leading causes of mortality worldwide. The central pathological features of atherosclerosis include macrophage infiltration and foam cell formation. However, the detailed mechanisms regulating these two processes remain unclear. Here we show that oxidative stress-activated Ca^2+^-permeable TRPM2 plays a key role in the pathogenesis of atherosclerosis. *Trpm2* deletion produces a potent protective effect against atherosclerosis in *ApoE*^-/-^ mice fed with a high-fat diet (HFD), as evidenced by reduced atherosclerotic plaque burden, decreased macrophage load and suppressed inflammasome activation in the vessel wall. Moreover, we show that *Trpm2* deletion or inhibition reduces oxidized low-density lipoprotein (oxLDL) uptake by macrophages, suppresses macrophage infiltration induced by monocyte chemoattractant protein-1 (MCP1), and prevents the impairment of macrophage emigration caused by oxLDL. Intriguingly, we uncover that activation of CD36, an oxLDL receptor, can promote the activation of TRPM2, and vice versa, the CD36-mediated inflammatory cascade in atherosclerosis is dependent on TRPM2. In transfected HEK293T cells, CD36 ligands oxLDL and TSP1 induce TRPM2 activation in a CD36-dependent manner. Deleting *Trpm2* or inhibiting TRPM2 activity in cultured macrophages suppresses the CD36 signaling cascade induced by oxLDL and TSP1. Our studies establish TRPM2-CD36 axis as a new mechanism underlying atherogenesis, and suggest TRPM2 as an effective therapeutic target for atherosclerosis.

**HIGHLIGHTS:**

- *Trpm2* deletion protects against atherosclerosis in *ApoE*^-/-^ mice fed with a high-fat diet (HFD)
- *Trpm2* deficiency reduces atherosclerotic lesions by minimizing foam cell formation, inhibiting macrophage infiltration and preserving macrophage emigration
- TRPM2 activation is required for CD36-induced oxLDL uptake and subsequent inflammatory responses
- The ligands of CD36, oxLDL and TSP1, activate TRPM2, thereby perpetuating TRPM2-CD36 inflammatory cycle in atherogenesis cascade
- Our data establish TRPM2-CD36 axis as a new atherogenesis mechanism and TRPM2 as a novel therapeutic target for atherosclerosis

TRPM2-mediated Ca^2+^ signal is essential for CD36 induced oxLDL uptake and atherosclerosis in *ApoE*^-/-^ mice fed with a high-fat diet (HFD). The activation of CD36 and TRPM2 form a positive feedback loop in atherogenesis.

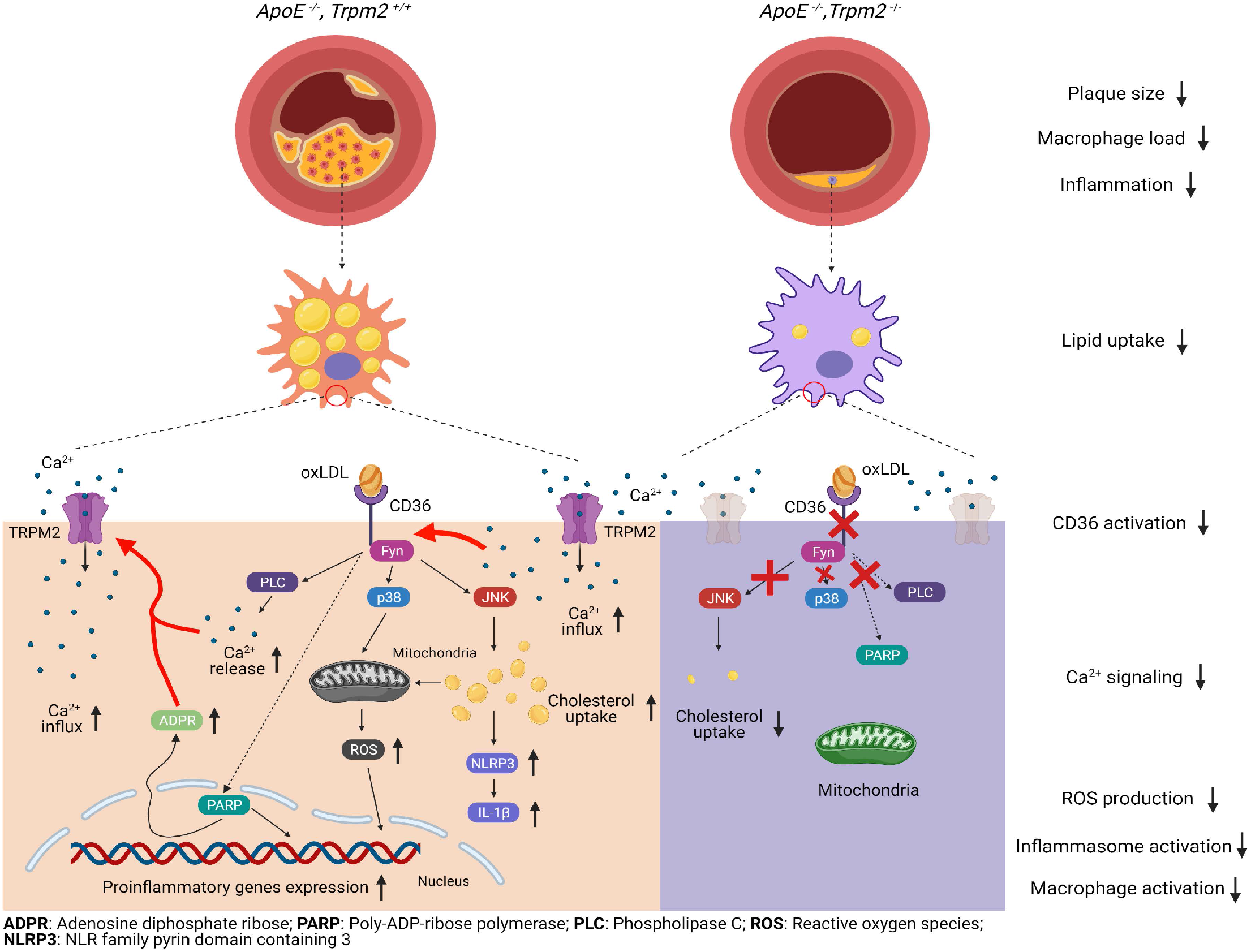

## INTRODUCTION

Atherosclerosis and its complications, such as myocardial infarction and stroke, are the leading cause of death worldwide^1^. Atherosclerosis is considered a chronic inflammatory disease of vessel wall. The initial and central pathological feature of atherosclerosis is the formation of foam cells after infiltrated macrophages phagocytize oxidized low density lipoprotein (oxLDL) and become overloaded with cholesterol^2^. These lipid-laden macrophages are the culprit for the progression of atherosclerotic lesions by secreting pro-inflammatory cytokines and matrix-degrading proteases, which cause profound inflammatory responses and tissue damage in vessel wall^3^. Therefore, inhibiting foam cell formation and inflammatory cytokine production could be a promising target for developing more effective therapies for atherosclerosis^1^.

The phagocytosis of oxLDL by macrophages is mediated by several scavenger receptors. CD36 is the most predominant oxLDL receptor as it is responsible for over 70% of oxLDL uptake^4^. The binding of oxLDL to CD36 not only triggers internalization of cholesterol, but also elicits downstream signaling cascades, including Fyn, JNK and p38, which further induces oxidative stress and expression of pro-inflammatory genes^5^. Moreover, binding of oxLDL to CD36 promotes the activation of NLRP3 inflammasome by interacting with Toll like receptor 4 and 6 (TLR4/6) heterodimer, driving the differentiation of macrophages toward a pro-inflammatory phenotype^6^. Thus, CD36 plays a critical role in the activation of macrophages and formation of foam cells in atherosclerotic lesions. However, the underlying mechanisms regarding how oxLDL binding triggers the activation of the CD36 signaling cascade in atherogenesis remain unclear^4,7^.

TRPM2 is a nonselective cation channel activated by reactive oxygen species (ROS), intracellular Ca^2+^, and ADP-ribose (ADPR)^8–10^, which are substantially generated in inflammatory responses^11^. TRPM2 is widely expressed in myeloid cells, and TRPM2 mediated Ca^2+^ signaling is important for macrophage activation and phagocytic functions^12,13^. Knockout of *Trpm2* was found to reduce the production of ROS in macrophages and mitigate tissue damage in a lung injury mouse model^14^. However, whether TRPM2 is involved in foam cell formation and atherogenesis is unknown. Considering atherosclerosis is also an inflammatory disease and TRPM2 is activated by oxidative stress under inflammatory conditions, we proposed that TRPM2 plays a key role in atherogenesis by integrating extracellular stimuli and intracellular signaling cascade.

In this study, we demonstrate that *Trpm2* deletion protects *ApoE*^-/-^ mice against HFD-induced atherosclerosis, characterized by reduced atherosclerotic lesions, decreased macrophage burden, and suppressed inflammasome activation in the vessel wall. We find that deletion of *Trpm2* or inhibiting the activation of TRPM2 in macrophages reduces oxLDL uptake, inhibits macrophage infiltration, and improves the impaired macrophage emigration. We reveal that the mechanism by which *Trpm2* deletion inhibits macrophage uptake of oxLDL is mediated by reduced CD36 activity. Moreover, we demonstrate that TRPM2 can be activated by the CD36 ligands in a CD36 dependent manner. Our studies establish a novel, mutually regulating, and positive feedback mechanism between CD36 and TRPM2 in atherogenesis. Targeting TRPM2 inhibits both TRPM2-dependent and CD36-dependent inflammatory response thereby producing strong protective effects against atherosclerosis.

## RESULTS

### *Trpm2* deletion protects *ApoE*^-/-^ mice from high-fat diet induced atherosclerosis

To investigate whether TRPM2 plays a role in atherosclerosis, *Trpm2*^-/-^ mice were crossed with *ApoE*^-/-^ mice. Successful *Trpm2* deletion was confirmed by PCR (Supplementary Fig. 1a), Western blot (Supplementary Fig. 1b) and whole-cell current recordings (Supplementary Fig. 1c-e). After mice were fed with a high-fat food (HFD) for 4 months, *ApoE*^-/-^ mice developed severe atherosclerosis, with a lesion ratio of 0.36±0.17. In contrast, *Trpm2* and *ApoE* double knockout mice exhibited significantly reduced atherosclerotic plaque lesion ratio (0.13±0.02) compared with *ApoE* single knockout (Fig. 1a,b), indicating that *Trpm2* deletion protects mice against atherogenesis.

**Figure 1:**
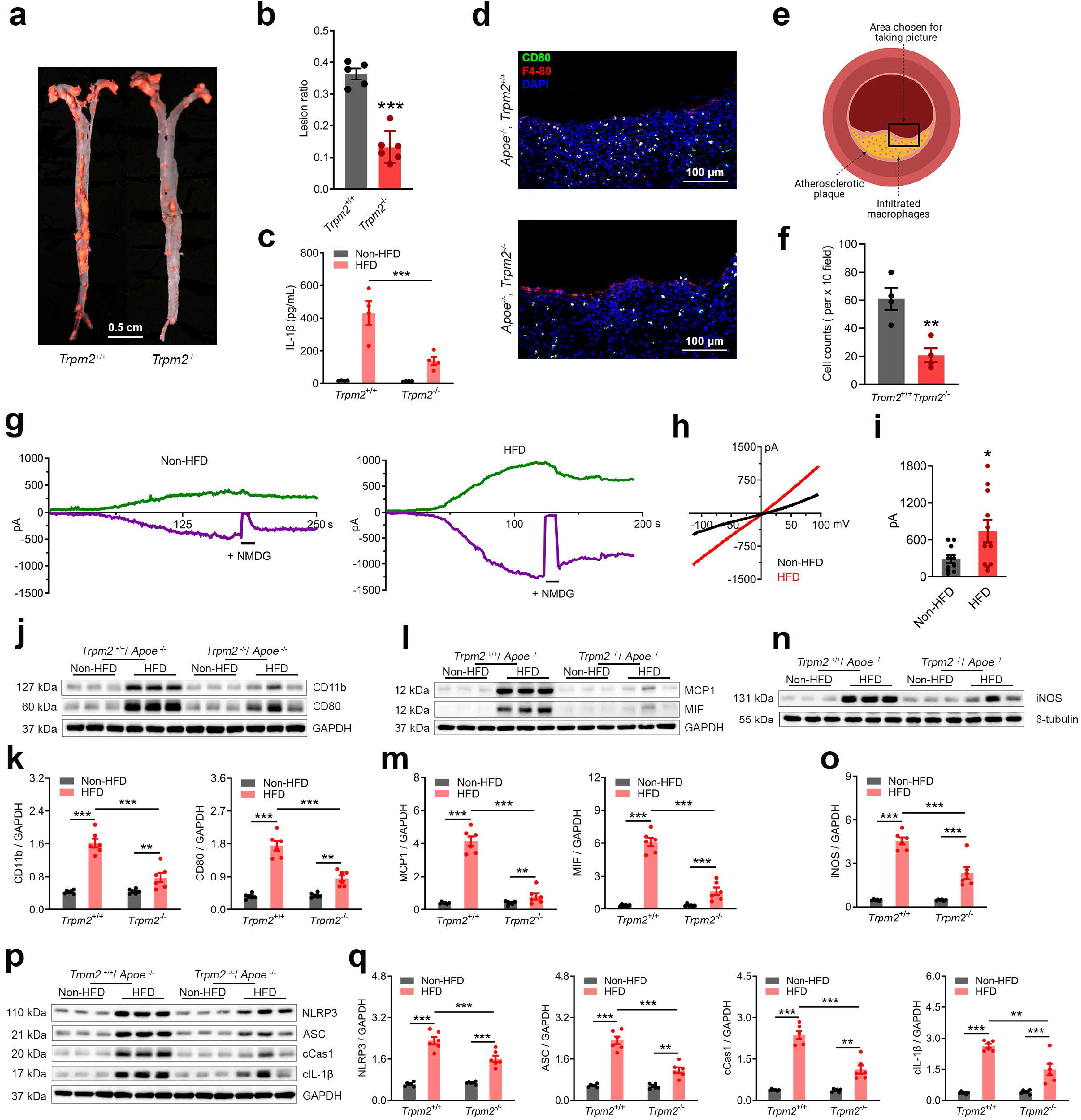
*Trpm2* deletion protects *ApoE*^-/-^ mice from high-fat diet induced atherosclerosis. (**a,b**) *Trpm2* deletion (*Trpm2*^-/-^) inhibited the formation of atherosclerotic plaque. **a**, Representative images of Oil Red O (ORO) staining of full-length aorta. **b**, Mean atherosclerotic lesion ratio based on ORO staining from *Trpm2*^+/+^ (n=6) and *Trpm2*^-/-^ mice (n=5). (**c**) *Trpm2* deletion (*Trpm2*^-/-^) inhibited systemic inflammation. Measurement of IL-1β level in serum from *Trpm2*^+/+^ and *Trpm2*^-/-^ mice with *ApoE*^-/-^ background using ELISA. (**d-f**) *Trpm2* deletion (*Trpm2*^-/-^) reduced macrophage burden in atherosclerotic plaque. **d**, Representative merged images of F4/80 and CD80 staining of aorta sections (Red: F4/80; Blue: DAPI; Green: CD80). e, Graphic illustration showing the atherosclerotic area chosen for taking pictures in **d** and **f. f**, Quantification of F4/80 and CD80 positive macrophages. 4 mice from each group were chosen for quantification. (**g-i**) **g**, Representative TRPM2 current traces (Green: outward current at +100 mV; Purple: inward current at +100 mV) in isolated peritoneal macrophages. NMDG blocks inward current indicating the tightness of seal. **h**, I-V relationship of TRPM2 current. **i**, Quantification of current amplitude (**j, k**) Representative western blot analysis and quantification of the expression of CD11b and CD80 in aorta. 6 mice from each group were chosen for quantification. (**l, m**) Representative western blot analysis and quantification of the expression of MCP1 and MIF in aorta. 6 mice from each group were chosen for quantification. (**n, o**) Representative western blot analysis and quantification of iNOS expression in aortas. 6 mice from each group were chosen for quantification. (**p, q**) Representative Western blot analysis and quantification of the expression of NLRP3, ASC, cleaved caspase-1 (cCAS1), and cleaved IL-1β (cIL-1β) expression in aortas. 6 mice from each group were chosen for quantification. (**: p < 0.01; ***: p < 0.001; ANOVA, Bonferroni’s test; mean ± SEM).

Atherosclerosis is a chronic inflammatory disease and is associated with systemic inflammation^15,16^. Indeed, in *ApoE* single knockout mice fed with HFD for 4 months, there was a dramatic increase of interleukin-1β (IL-1β) level in serum compared to mice fed with regular chow (430.70±73.69 pg/mL), whereas this increase was significantly attenuated in *Trpm2* and *ApoE* double knockout mice (138.03±27.10 pg/mL) (Fig. 1c). This decrease of circulating IL-1β might result from the alleviated atherosclerotic lesion on aorta, and indicates that global *Trpm2* deletion attenuates systemic inflammation caused by HFD treatment in *ApoE*^-/-^ mice. Consistent with the role of TRPM2 in systemic inflammation, TRPM2 current amplitude recorded in peritoneal macrophages isolated from HFD fed mice was significantly larger than that from regular chow fed mice (Fig. 1g-i).

Macrophage infiltration plays a critical role in the initiation and progression of atherosclerosis^3^. To understand whether TRPM2 plays a role in macrophage infiltration, we evaluated M1 macrophages, a pro-inflammatory subtype, in the atherosclerotic vessel by using immunostaining with anti-CD80. We found that the number of F4/80 and CD80 positive macrophages in atherosclerotic plaque was reduced from 61.00±7.80 per x 10 field in *ApoE* single knockout mice to 20.75±5.11 per x 10 field in *Trpm2* and *ApoE* double-knockout mice (Fig. 1d-f), indicating that macrophage burden is significantly reduced by *Trpm2* deletion. Consistent with the reduced macrophage numbers, the expression level of CD11b, another surface marker of macrophages, as well as CD80 assessed by Western blot, were markedly lower in the aorta of *Trpm2* and *ApoE* double-knockout mice than *ApoE* single-knockout mice after 4-month HFD treatment (Fig. 1j,k). These data indicate that *Trpm2* deletion inhibits the increase of macrophage burden in aorta during atherogenesis.

Macrophage infiltration in atherosclerosis is mainly influenced by two chemokines, monocyte chemoattractant protein-1 (MCP1) and macrophage migration inhibitory factor (MIF)^17^. At the initial stage of atherosclerosis, MCP1 secreted by endothelial cells upon subendothelial oxLDL deposition is a major cause of macrophage infiltration^3^, whereas during the progression of atherosclerosis, MCP1 and MIF secreted by activated macrophages themselves and smooth muscle cells further aggregate the infiltration of macrophages^17–20^. Moreover, MCP1 and MIF promotes the differentiation of macrophages toward a pro-inflammatory phenotype^20,21^. To understand the mechanisms by which *Trpm2* deletion reduces macrophage burden, we analyzed the expression levels of MCP1 and MIF in the atherosclerotic aorta by WB. *Trpm2* deletion drastically reduced both MCP1 and MIP levels in the aorta of *ApoE*^-/-^ mice fed with a HFD for 4 months (Fig. 1l,m). These data strongly suggests that *Trpm2* deletion reduces the number of macrophages in the lesion area (Fig. 1d) by inhibiting MCP1 and MIP expression.

Macrophages that infiltrate into the atheroprone site quickly become the center of inflammatory cues. Inducible nitric oxide synthase (iNOS) was previously found to be abundantly expressed in human atherosclerotic lesions^22^. Since iNOS shifts the production of nitric oxide by NOS toward the production of ROS and promotes the activation of macrophages^23^. Therefore, we evaluated whether iNOS expression is influenced by TRPM2. We found that *Trpm2* deletion significantly inhibited the increase of iNOS expression in aorta after HFD treatment (Fig. 1n,o).

Activation of NLRP3 inflammasome by phagocytized cholesterol crystals in macrophages is required for atherogenesis^24^. We therefore determined whether TRPM2 is involved in inflammasome activation. Our result showed that in *ApoE* single knockout mice fed a HFD for 4 months, there was a significant increase of NLRP3, ASC, cleaved Caspase1 and cleaved IL-1β expression in the aorta compared to the mice fed with regular chow, whereas this increase was attenuated in *Trpm2* and *ApoE* double knockout mice (Fig. 1p, q). In summary, the above results indicate that *Trpm2* deletion reduces macrophage infiltration and mitigates inflammation in aorta induced by HFD treatment.

### Deletion of *Trpm2* reduces the uptake of oxLDL by macrophages, suppresses macrophage infiltration, preserves macrophage emigration and inhibits macrophage activation *in vitro*

Macrophage infiltration is a critical step in initiating atherogenesis^3^. We therefore established a new *in vitro* assay to examine whether TRPM2 influences the infiltration ability of macrophages (Fig. 2a). Different from previous reported infiltration assays^25^, aortic endothelial cells isolated from wild-type (WT) mice were plated onto the upper surface of transwell inserts to better simulate the pathophysiological conditions during atherosclerosis. After endothelial cells completely covered the upper surface of inserts with 12 μm pore size, bone marrow derived macrophages isolated from either WT or *Trpm2* knockout (M2KO) mice were added into the upper chamber, while MCP1 was added into the lower chamber to promote macrophage infiltration. 24 h after adding macrophages into the upper chamber, the 25 mm cover slips on the bottom of the lower chamber were collected for detecting infiltrated macrophages using F4/80 and CD80 co-staining (Fig. 2b). We found that MCP1 induced a significant increase of macrophage infiltration into the lower chamber in WT compared to PBS, but this increase was inhibited in M2KO group (Fig. 2b,c), indicating that *Trpm2* deletion reduces the infiltration ability of macrophages.

**Figure 2:**
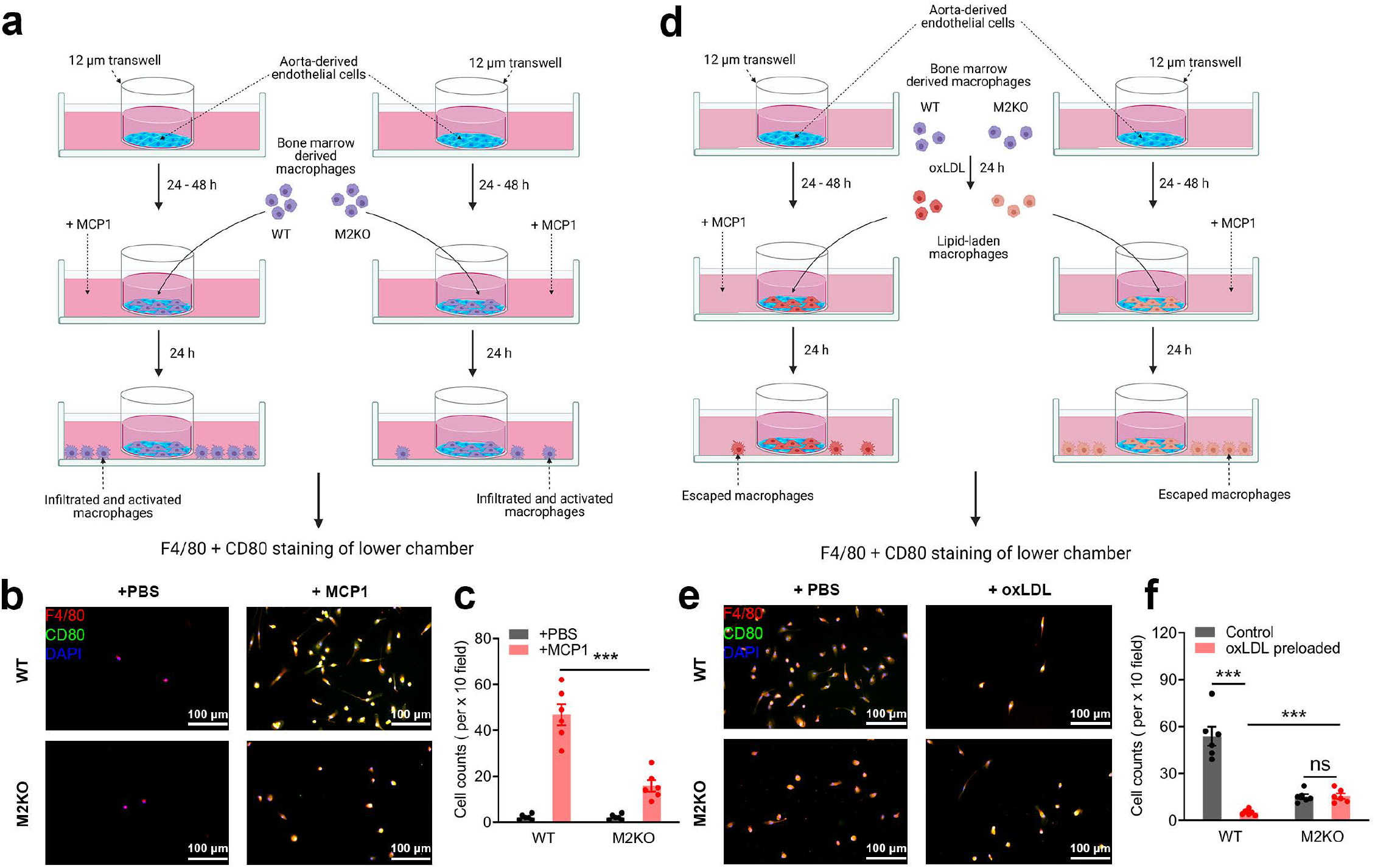
Deletion of *Trpm2* inhibits macrophage infiltration while preserves macrophage emigration. (**a-c**) *Trpm2* deletion inhibits macrophage infiltration. **a**, Graphic illustration of *in vitro* examination of macrophage infiltration across endothelial cells induced by MCP1. Aorta-derived endothelial cells were plated on the transwell inserts (pore size: 12 μm) for 3-5 days. Bone marrow derived macrophages were added into the upper chamber after endothelial cells completely covered the upper surface of transwells. After 24 h, F4/80 and CD80 staining of macrophages in the lower chamber was performed as in **b** (Red: F4/80; Blue: DAPI; Green: CD80). **c**, Quantification of the number of infiltrated macrophages within a x 10 field. 6 dishes from each group were chosen for quantification (**d-f**) *Trpm2* deletion prevented the loss of emigration ability in oxLDL-pre-loaded macrophages. **e**, Graphic illustration of *in vitro* examination of macrophage emigration across endothelial cells induced by MCP1. Aorta-derived endothelial cells were plated on the transwell inserts (pore size: 12 μm) for 3-5 days. Bone marrow derived macrophages preloaded with oxLDL for 24 h were added into the upper chamber after endothelial cells completely covered the upper surface of transwells. After 24 h, F4/80 and CD80 staining of macrophages in lower chamber was performed as in **e** (Red: F4/80; Blue: DAPI; Green: CD80). **f**, Quantification of the number of infiltrated macrophages with in a x 10 field. 6 dishes from each group were chosen for quantification. (ns: no statistical significance; ***: p < 0.001; ANOVA, Bonferroni’s test; mean ± SEM).

Macrophage emigration refers to the returning of macrophages from atherosclerotic lesion sites back into circulation, which is important for atherosclerosis plaque regression^2,19^. However, phagocytizing oxLDL markedly impairs the migration ability of macrophages, resulting in macrophages being trapped in atherosclerotic areas thereby sustaining inflammatory responses in the vessel wall^25^. To investigate the role of TRPM2 in macrophage emigration, we designed a new *in vitro* assay to examine the emigration ability of macrophages (Fig. 2d). All the steps are the same to the infiltration test shown in Fig. 2a, except that macrophages were preloaded with oxLDL for 24 h (Fig. 2d). We found that preloading with oxLDL dramatically inhibited the migration of macrophages in WT cells, but not in M2KO cells (Fig. 2e,f). Our results indicate that *Trpm2* deletion not only inhibits macrophage infiltration induced by MCP1 (Fig. 2a), but also eliminates the impairment of macrophage emigration caused by oxLDL (Fig. 2b).

Over-phagocytosis of oxLDL transforms macrophages into highly pro-inflammatory foam cells and thereby inhibits macrophage emigration^2^. To determine whether TRPM2 influences oxLDL engulfment, we used Oil Red O staining to evaluate the formation of foam cells as previously reported^26^. We found that deletion of *Trpm2* markedly inhibited the uptake of oxLDL in macrophages (Fig. 3a,b). As NLRP3 inflammasome activation induced by uptake of oxLDL is critical for macrophage activation during atherogenesis^6^, we measured the oxLDL-induced production of IL-1β and found that *Trpm2* deletion inhibited IL-1β secretion induced by oxLDL, indicating that activation of NLRP3 inflammasome was suppressed by *Trpm2* deletion (Fig. 3c).

**Figure 3:**
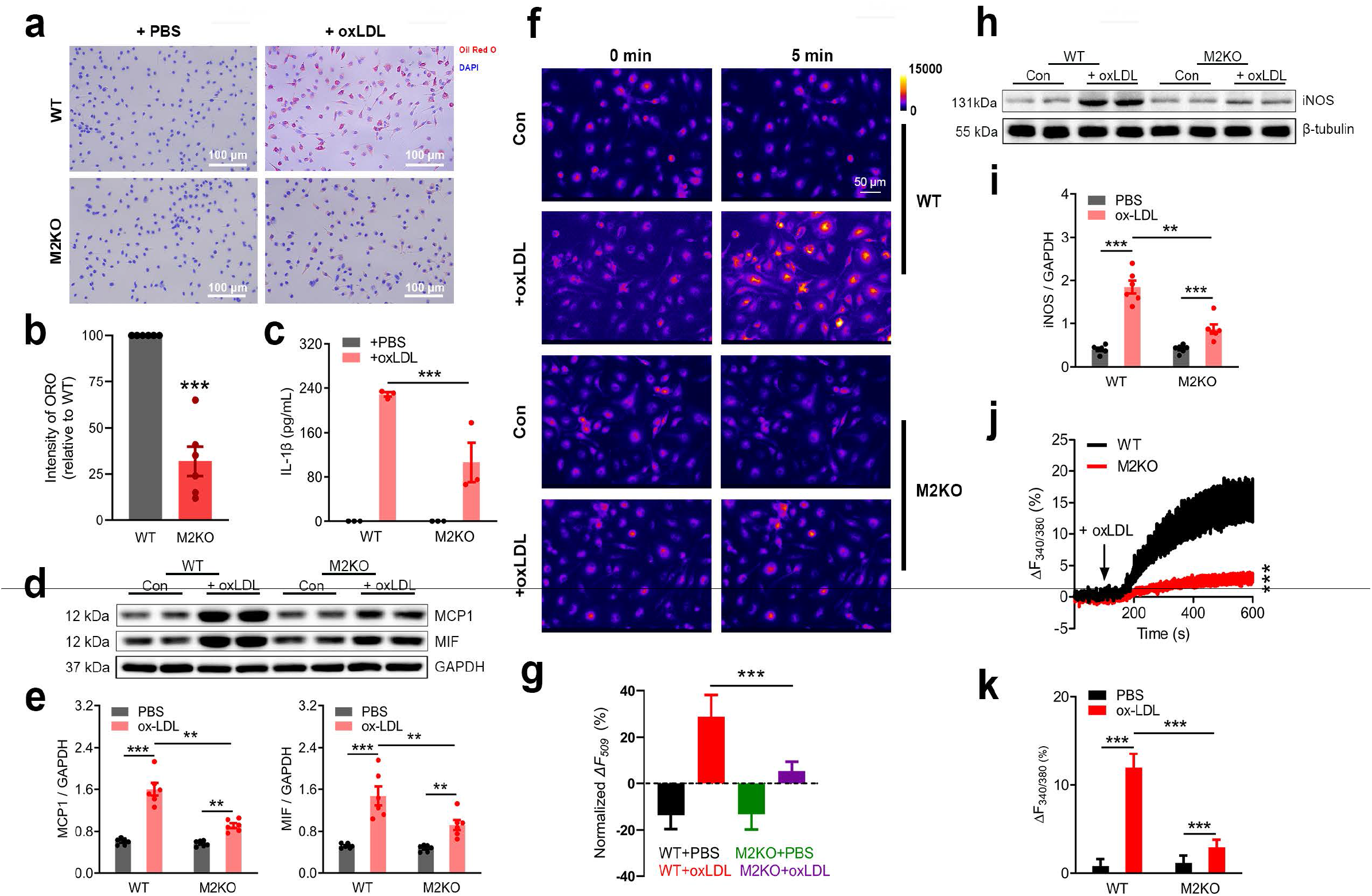
Deletion of *Trpm2* inhibits the uptake of oxLDL and pro-inflammatory activation of macrophages. (**a, b**) Representative images and quantification of Oil Red O (ORO) staining in isolated macrophages from wild-type (WT) or *Trpm2* knockout (M2KO) mice after treatment with oxLDL (50 μg/ml) for 24 h. 6 dishes of cells from 6 mice from each group were chosen for quantification. (**c**) *Trpm2* deletion inhibited the production of IL-1β in macrophages. Measurement of IL-1β level in culture medium after the treatment of oxLDL (50 μg/ml) for 24 h using ELISA. 3 dishes of cells from 3 mice from each group were chosen for quantification. (**d, e**) Representative western blot analysis and quantification of the expression of MCP1 and MIF in isolated macrophages after oxLDL treatment (50 μg/ml) for 24 h. 6 dishes of cells from 6 mice from each group were chosen for quantification. (**f, g**) **f**, Representative picture of Rhodamine-123 real-time imaging before and 5 min after oxLDL treatment (50 μg/ml) in isolated macrophages. Control group (PBS treatment) was used to show the rapid photo bleaching of R123. **g**, Quantification of changes of R123 fluorescence 5 min after oxLDL treatment. WT (n=40 for oxLDL treatment, n=38 for control) and M2KO (n=43 for oxLDL treatment, n=40 for control) macrophages were from 4 dishes of cultured cells isolated from 4 mice in each group. (**h, i**) Representative Western blot analysis and quantification of the expression of iNOS in isolated macrophages treated with oxLDL (50 μg/ml). 6 dishes of cells from 6 mice from each group were chosen for quantification (**j, k**) **j**, Representative real-time Fura-2 Ca^2+^ imaging traces during oxLDL treatment (50 μg/ml). The averaged traces were from 10 macrophages randomly chosen from a representative culture dish for each groups. **k**, Quantification of Fura-2 fluorescence changes 5 min after oxLDL treatment. WT (n=20 for oxLDL treatment, n=20 for control) and M2KO (n=20 for oxLDL treatment, n=20 for control) macrophages were from 3 dishes of cultured cells isolated from 3 mice in each group. (**: p < 0.01; ***: p < 0.001; ANOVA, Bonferroni’s test; mean ± SEM).

In atherosclerotic plaque, persistent inflammation promotes activated macrophages to secrete MCP1 and MIF, which lead to the recruitment of more macrophages into the lesion site, thereby resulting in a positive feed-back vicious cycle that accelerates atherosclerosis progression^3^. We therefore analyzed whether TRPM2 influences MCP1 and MIF production. Western blot analysis revealed that the expression of MCP1 and MIF were significantly increased after oxLDL treatment in WT macrophages, but not in M2KO macrophages (Fig. 3d,e), indicating that deletion of *Trpm2* results in reduced MCP1 and MIF production by macrophages. It is conceivable that the decreased levels of MCP1 and MIF in *Trpm2* deletion mice result in reduced infiltration and the preserved emigration of macrophages, thereby leading to reduced macrophage burden in aorta as shown in Fig. 1e.

In macrophages, digestion of oxLDL leads to the substantial production of reactive oxygen species (ROS), which activates signaling pathways such as nuclear factor-κB (NF-κB) pathway, a key signaling cascade in activating inflammation related genes^23,27^. Rhodamine-123 (R123) imaging is a commonly used method to monitor mitochondria oxidative stress and ROS production^28,29^. Using real-time recording of R123 fluorescence, we found that oxLDL induced a marked and rapid increase of R123 signal in WT but not M2KO macrophages within 5 min of oxLDL exposure (Fig. 3f,g). To understand how TRPM2 influences ROS production, we evaluated the level of iNOS, a known factor that promotes the production of ROS in macrophages^23^. As shown in Fig. 3h,i, *Trpm2* deletion significantly inhibited the increase of iNOS expression in macrophages induced by oxLDL.

Increasd intracellular Ca^2+^ is crucial for macrophage activation^12,30^. We therefore examined changes of Ca^2+^ signaling in response to oxLDL stimulation using Fura-2 real-time Ca^2+^ imaging. We found that oxLDL induced a robust increase of intracellular Ca^2+^ concentration in WT macrophages, but this increase was significantly inhibited in M2KO macrophages (Fig. 3j,k), suggesting that oxLDL-induced intracellular Ca^2+^ changes are dependent on TRPM2. As both ROS and Ca^2+^ signaling are critical for macrophage activation in response to inflammatory stimulation^13,23,27^, the inhibitory effects of *Trpm2* deletion on ROS production and intracellular Ca^2+^ signaling further indicate that knockout of *Trpm2* inhibits oxLDL-mediated activation of macrophages.

### *Trpm2* deletion impairs the activation of CD36 signaling cascades by oxLDL and thrombospondin 1 (TSP1)

The strong inhibition of oxLDL uptake as well as its downstream signaling pathways by *Trpm2* deletion prompted us to investigate the mechanism by which TRPM2 influences oxLDL uptake in atherogenesis. CD36 is the major receptor mediating oxLDL uptake, and is responsible for over 70% of oxLDL uptake in macrophages^5^. Activating signaling cascades downstream of CD36, such as Fyn, JNK, and p38 have been shown to promote the activation of macrophages, and inhibit the emigration of macrophages from atherosclerotic plaques^5,25^. Activation of p38 MAP kinase and JNK2 are required for foam cell formation^31,32^, and CD36-mediated activation of JNK2 is necessary for oxLDL uptake^33^. Based on our observation of reduced oxLDL uptake in macrophages and inhibited macrophage activation by *Trpm2* deletion, we proposed that TRPM2 influences oxLDL-induced atherogenesis by regulating CD36 function.

Indeed, we found that the oxLDL treatment-induced upregulation of CD36 as well as increased phosphorylation of Fyn, JNK and p38 in WT macrophages were largely minimized in M2KO macrophages (Fig. 4a,b). To exclude the non-specific effect caused by oxLDL and confirm the role of TRPM2 in activating CD36 signaling cascades, we used another CD36 ligand TSP1. TSP1 is an extracellular glycoprotein secreted by macrophages and other types of cells, and is known to promote inflammation^34,35^. TSP1 was found to activate macrophages and promote the production of tumor necrosis factor α in macrophages by activating the NF-κB pathway in a CD36-dependent manner^36^. It was recently demonstrated that deletion of TSP1 protects mice against leptin induced atherosclerosis by inhibiting the abnormal activation of smooth muscle cells^37^. However, whether TSP1 is involved in HFD induced atherogenesis remains unknown. We found that similar to oxLDL, TSP1 elicited upregulation of CD36 and increased phosphorylation of Fyn, JNK, and p38 in WT macrophages but not in M2KO macrophages (Fig 4c,d), indicating that TPRM2 is critical for the activation of the CD36 signaling cascade.

**Figure 4:**
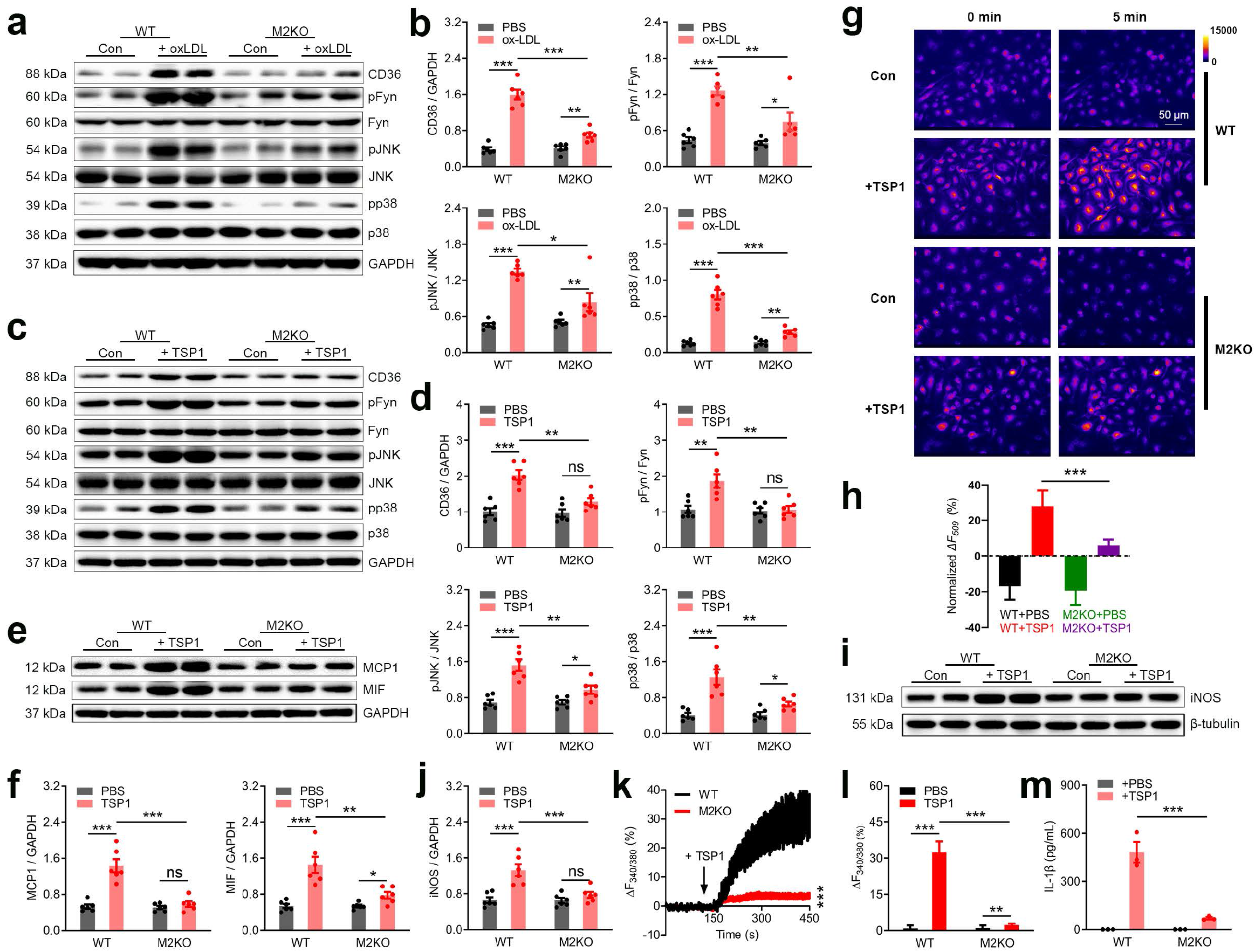
*Trpm2* deletion inhibits the activation of CD36 signaling cascades by oxLDL and TSP1. (**a-d**) **a**, **c** Representative western blot analysis of CD36, pFyn, Fyn, pJNK, JNK, pp38 and p38 expression in isolated macrophages after oxLDL (50 μg/ml) or TSP1 (10 μg/ml) treatment. **b**, **d** Quantification of western blot bands. 6 dishes of macrophages from 6 mice were used for protein extraction in each group. (**e, f**) Representative western blot analysis and quantification of the expression of MCP1 and MIF in isolated macrophages after treatment with TSP1 (10 μg/ml) for 24 h. 6 dishes of cells from 6 mice from each group were chosen for quantification. (**g, h**) **g**, Representative picture of R123 imaging before and 5 min after TSP1 treatment (10 μg/ml) in isolated macrophages. Control group (Con: PBS treatment) was used to show the rapid photo bleaching of R123. **h**, Quantification of changes of R123 fluorescence 5 min after TSP1 treatment. WT (n=45 for TSP1 treatment, n=47 for control) and M2KO (n=43 for TSP1 treatment, n=47 for control) macrophages were from 4 dishes of cultured cells isolated from 4 mice in each group. (**i, j**) Representative western blot analysis and quantification of iNOS expression in isolated macrophages treated with TSP1 (10 μg/ml). 6 dishes of cells from 6 mice from each group were chosen for quantification. (**k**), Representative real-time Fura-2 Ca^2+^ imaging traces during TSP1 treatment (10 μg/ml). The averaged traces were from 10 macrophages randomly chosen from a representative culture dish of each group. (**l**), Quantification of Fura-2 fluorescence changes 5 min after TSP1 treatment. WT (n=20 for TSP1 treatment, n=20 for control) and M2KO (n=20 for TSP1 treatment, n=20 for control) macrophages were from 3 dishes of cultured cells isolated from 3 mice in each group. (**m**) *Trpm2* deletion (M2KO) inhibited the production of IL-1β in macrophages. Measurement of IL-1β level in culture medium after the treatment of TSP1 (10 μg/ml) for 24 h using ELISA. 3 dishes of cells from 3 mice from each group were chosen for quantification. (ns: no statistical significance; *: p < 0.05; **: p < 0.01; ***: p < 0.001; ANOVA, Bonferroni’s test; mean ± SEM).

Next, we sought to determine the effects of TRPM2 on TSP1-induced pro-inflammatory cytokine secretion in macrophages. Fig. 4e,f show that TSP1 induced an increase in MCP1 and MIF1 expression in WT macrophages, but not in M2KO macrophages. Moreover, TSP1 treatment induced an increase of R123 fluorescence in WT macrophages but this increase was markedly attenuated in M2KO macrophages (Fig. 4g,h), suggesting that ROS production induced by TSP1 is inhibited by *Trpm2* deletion. Consistent with the reduced ROS production in M2KO macrophages, the expression of iNOS was also much lower in M2KO than WT macrophages (Fig. 4i,j), presumably due to reduced oxLDL uptake through CD36 in M2KO macrophages. These results indicate that *Trpm2* deletion inhibits TSP1-induced CD36 activation.

We next tested the effect of TSP1 on Ca^2+^ signaling. Interestingly, TSP1 at a lower concentration induced a larger increase of intracellular Ca^2+^ concentration than that induced by oxLDL (TSP1 at 10 μg/ml vs oxLDL at 50 μg/ml) in macrophages from WT group but not in M2KO group (Fig. 4k,l). Moreover, TSP1 treatment increased the secretion of IL-1β by macrophages, a marker representing the activation of NLRP3 inflammasome, whereas this increase was inhibited in M2KO macrophages (Fig. 4m). As the expression of TSP1 was shown to be significantly increased in the aorta with atherosclerotic lesions^38^, our data suggest that TSP1 induced activation of CD36 signaling plays an important role in macrophage activation in atherosclerotic plaques.

In summary, the above results suggest that both oxLDL and TSP1 activate CD36 in atherosclerosis, and that deletion of *Trpm2* inhibits oxLDL and TSP1 induced activation of CD36, indicating that TRPM2 is required for CD36 activation.

### TRPM2 mediates the activation of CD36 signaling cascades in macrophages induced by oxLDL and TSP1

The requirement of TRPM2 for CD36 activation by oxLDL and TSP1 is an interesting discovery. To understand the underlying mechanisms, we first determined whether activation of CD36 influences TRPM2 channel function. We confirmed that TRPM2 can indeed be activated during oxLDL or TSP1 treatment. We found that under the recording conditions with 500 nM Ca^2+^ and 1 μM ADPR in the pipette solution, oxLDL substantially activated TRPM2 currents in HEK293T cells co-transfected with both TRPM2 and CD36 in 5 min (Fig. 5a). In contrast, there was no TRPM2 current activation in cells transfected with TRPM2 only, even after perfusion with oxLDL for 10 min (Fig. 5b). The currents activated by oxLDL display typical TRPM2 characteristics such as linear I-V relation (Supplementary Fig. 2a) and can be blocked by 30 μM N-(p-amylcinnamoyl) anthranilic acid (ACA). Moreover, preincubation with sulfosuccinimidyl oleate (SSO), a CD36 specific inhibitor, effectively abolished the activation of TRPM2 by oxLDL (2156.00 ± 342.00 pA vs 50.02 ± 10.65 pA) (Figure 5c, Supplementary Fig. 2a).

**Figure 5:**
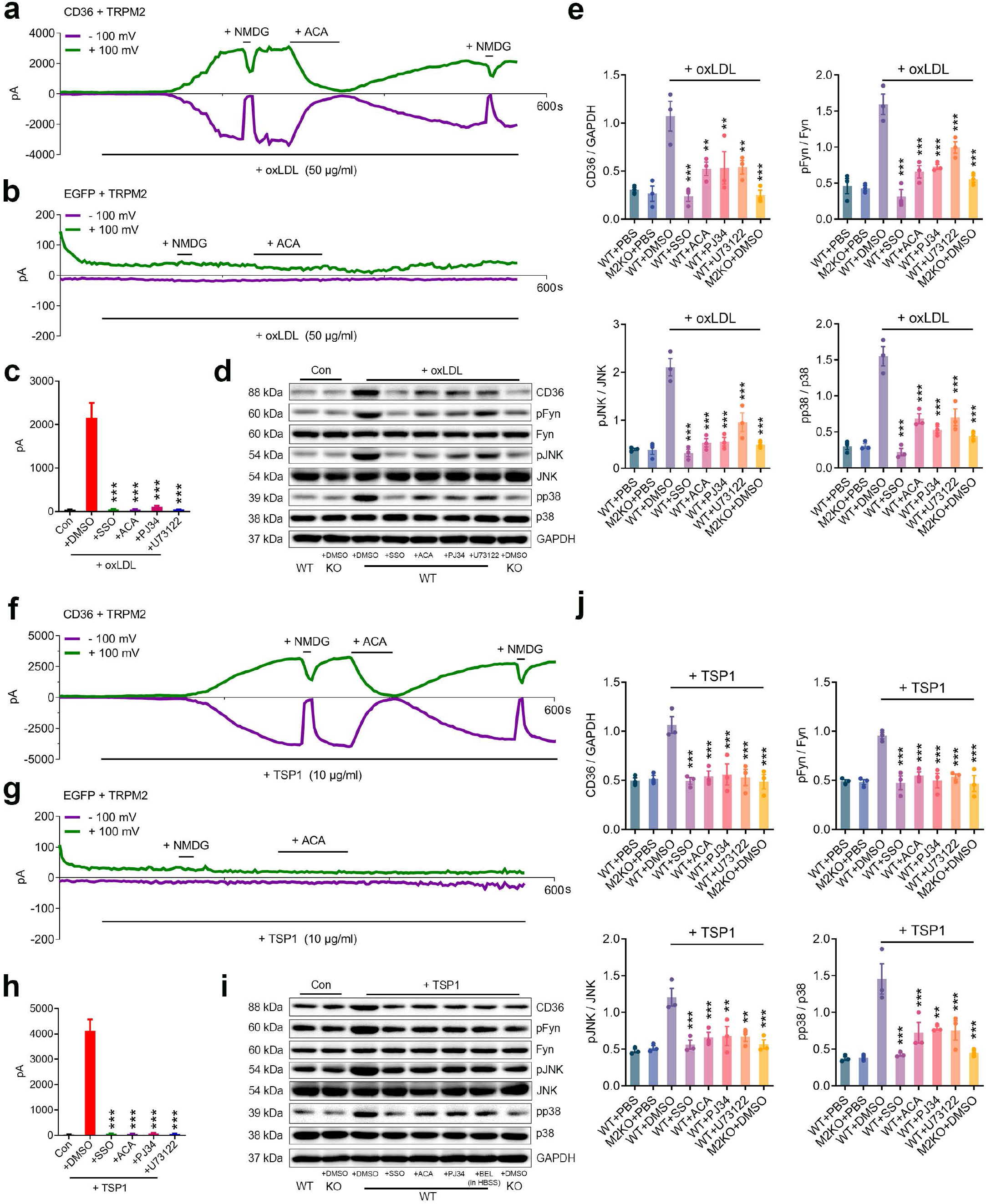
TRPM2 mediates the activation of CD36 signaling cascades in macrophages induced by oxLDL and TSP1. (**a-c**) CD36 is needed for the activation of TRPM2 induced by oxLDL (50 μg/ml) in HEK293T cells. **a**, Representative TRPM2 current traces (Green: outward current at +100 mV; Purple: inward current at −100 mV) in HEK293T cells transfected with CD36 and TRPM2 during oxLDL treatment. NMDG blocks inward current indicating the tightness of seal. ACA is a TRPM2 blocker. **b**, Representative recording traces in HEK293T cells transfected with only TRPM2 during oxLDL treatment. **c**, Quantification of TRPM2 current amplitude in HEK293T cells transfected with CD36 and TRPM2. (**d, e**) Inhibiting the activation of TRPM2 impairs the activation of CD36 signaling cascade induced by oxLDL (50 μg/ml) in macrophages. **d,** Representative western blot analysis of the expression of CD36, pFyn, Fyn, pJNK, JNK, pp38 and p38 in isolated macrophages from WT (n=3 in each group) and M2KO mice (n=3 in each group). **e**, Quantification of western blot bands. 3 dishes of macrophages were used for protein extraction in each group. (**f-h**) CD36 is needed for the activation of TRPM2 induced by TSP1 (10 μg/ml) in HEK293T cells. **f**, Representative TRPM2 current traces (Green: outward current at +100 mV; Purple: inward current at −+100 mV) in HEK293T cells transfected with CD36 and TRPM2 during TSP1 treatment. NMDG blocks inward current indicating the tightness of seal. ACA is a TRPM2 blocker. **g**, Representative recording traces in HEK293T cells transfected with only TRPM2 during TSP1 treatment. **h**, Quantification of TRPM2 current amplitude in HEK293T cells transfected with CD36 and TRPM2. (**i, j**) Inhibiting the activation of TRPM2 impairs the activation of CD36 signaling cascade induced by TSP1 (10 μg/ml) in macrophages. **i,** Representative Western blot analysis of the expression of CD36, pFyn, Fyn, pJNK, JNK, pp38 and p38 in isolated macrophages from WT (n=3 in each group) and eM2KO mice (n=3 in each group). **j**, Quantification of Western blot bands. 3 dishes of macrophages were used for protein extraction in each group. (*: p < 0.05; **: p < 0.01; ***: p < 0.001; ANOVA, Bonferroni’s test; mean ± SEM).

The activation of TRPM2 needs intracellular Ca^2+^ and ADP ribose (ADPR)^39^. Since our pipette solution only contains 500 nM Ca^2+^ and 1 μM ADPR, we reasoned that oxLDL might influence intracellular Ca^2+^ or ADPR thereby activating TRPM2. We found that PJ34, a specific inhibitor for poly ADP-ribose polymerase, abolished the activation of TRPM2 by oxLDL in HEK293 cells transfected with TRPM2 and CD36 during oxLDL treatment (Figure 5c, Supplementary Fig. 2a). Similarly, U73122, a potent PLC inhibitor, when used in combination with extracellular Ca^2+^ free recording solution (0.5 mM EDTA), abolished the activation of TRPM2 by oxLDL in HEK293 cells co-expressed with TRPM2 and CD36 during oxLDL treatment (Figure 5c, Supplementary Fig. 2a). Furthermore, inhibition of TRPM2 activation using ACA, PJ34, and U73122 (used in combination with Ca^2+^ free HBSS culture medium), significantly inhibited the activation of CD36 signaling cascades in macrophages during oxLDL treatment (Fig. 5d,e). TRPM2 is a non-selective Ca^2+^-permeable cation channel^40^. TRPM2 mediated Ca^2+^ signaling was found to be important for various cellular functions^40^. Given *Trpm2* deletion or inhibiting the activation of TRPM2 produced a similar inhibitory effect on CD36 signaling cascade as using Ca^2+^ free medium (HBSS), our result indicates that TRPM2 mediated Ca^2+^ signaling is important for oxLDL activated CD36 signaling cascades in macrophages.

Interestingly, compared to oxLDL, TSP1 treatment induced a more robust activation (4117.00 ± 454.90 pA by TSP1 versus 2156.00 ± 342.00 pA by oxLDL) of TRPM2 currents in HEK293T cells transfected with both TRPM2 and CD36 (Fig. 5f), and this activation did not happen in HEK293T cells transfected with TRPM2 alone (Fig. 5g). Preincubation with SSO completely inhibited the activation of TRPM2 by TSP1 (Figure 5h, Supplementary Fig. 2b). Similar to oxLDL treatment, the activation of TRPM2 by TSP1 disappeared when transfected cells were preincubated with ACA, PJ34 and U73122 (Figure 5h, Supplementary Fig. 2b). Also, the activation of CD36 signaling pathway in macrophages by TSP1 was significantly inhibited with ACA, PJ34, and U73122 (used in combination with Ca^2+^ free HBSS culture medium) treatment(Fig. 5i,j), likely by inhibiting TRPM2 activation. Moreover, SSO, ACA, PJ34 and U73122 treatments did not produce an additional effect on the activation of CD36 signaling cascades in macrophages subjected to oxLDL (Supplementary Fig. 3a,b) or TSP1 treatment (Supplementary Fig. 3c,d), indicating these inhibitors did not affect CD36 in the absence of TRPM2. The above data suggest that during oxLDL and TSP1 treatment, CD36 signaling activates TRPM2 by increasing the production of ADPR and intracellular Ca^2+^ concentration, and TRPM2 mediated Ca^2+^ signaling is also critical for the activation of CD36 signaling cascades.

### TRPM2 mediates ROS production, increased Ca^2+^ concentration and inflammasome activation in macrophages induced by oxLDL or TSP1

We then sought to understand whether inhibiting the activation of TRPM2 influences the activation of macrophages by oxLDL or TSP1. As a negative control, inhibition of CD36 by preincubating macrophages with SSO significantly mitigated the increase of R123 signal in macrophages induced by either oxLDL (Fig. 6a,c) or TSP1 treatment (Fig. 6b,d). Moreover, ACA, PJ34 and U73122 preincubation markedly inhibited the increase of R123 signal in macrophages induced by either oxLDL (Fig. 6a,c) or TSP1 treatment (Fig. 6b-d). Furthermore, TRPM2 inhibitor ACA and inhibiting the activation of TRPM2 by PJ34 or U73122 (with Ca^2+^ free extracellular medium) inhibited the expression iNOS in macrophages receiving oxLDL (Fig. 6e,g) and TSP1 treatment (Fig. 6f,h). Notably, SSO, ACA, PJ34 and U73122 treatments did not additionally inhibit the expression of iNOS in M2KO macrophages subjected to oxLDL (Supplementary Fig. 3e,f) or TSP1 treatment (Supplementary Fig. 3g,h). Similarly, SSO, ACA, PJ34 and U73122 preincubation markedly inhibited the increase of intracellular Ca^2+^ concentration in macrophages induced by either oxLDL (Fig. 6i,k) or TSP1 treatment (Fig. 6j-l). These data indicate that inhibiting the activation of TRPM2 suppressed the production of ROS and the increase of intracellular Ca^2+^ concentration in macrophages during oxLDL or TSP1 treatment. Since ROS and Ca^2+^ signaling are critical for the activation of pro-inflammatory pathways in macrophages^12,27^, we examined whether these inhibitors affect the activation of NLRP3 inflammasome by measuring the concentration of IL-1β in culture medium. Our data showed that by inhibiting the activation of TRPM2 using ACA, PJ34 and U73122, the secretion of IL-1β by macrophages induced by oxLDL or TSP1 was significantly inhibited (Fig. 6m,n). Considering the crucial role of ROS and Ca^2+^ signaling in the activation of macrophages, the above results suggest that inhibition of TRPM2 activation significantly suppresses macrophage activation induced by oxLDL and TSP1.

**Figure 6:**
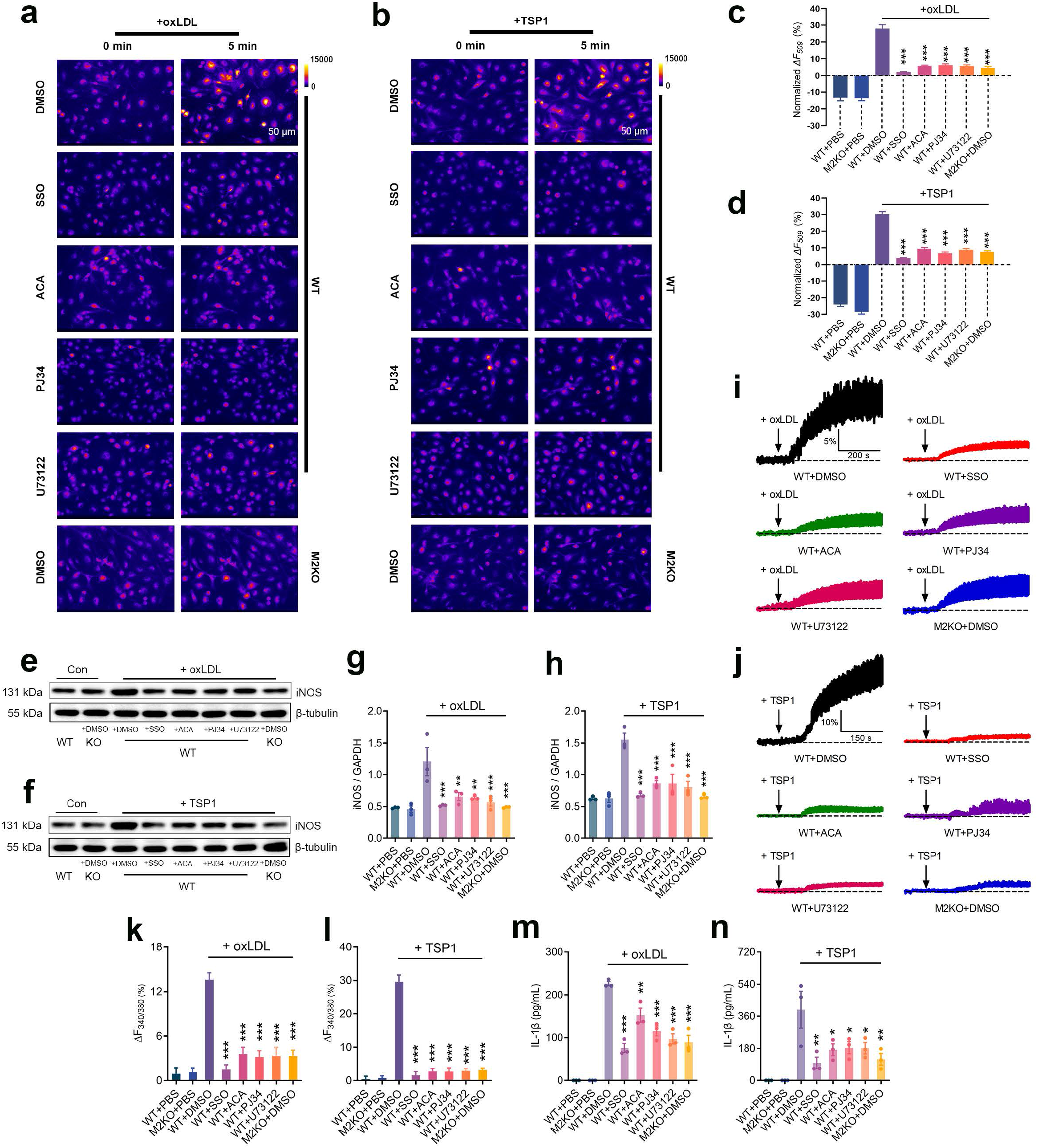
TRPM2 mediates ROS production, increased Ca^2+^ concentration and inflammasome activation in macrophages induced by oxLDL or TSP1. (**a-d**) Representative picture of Rhodamine-123 real-time imaging of macrophages before and 5 min after oxLDL treatment (50 μg/ml) as in **a**, and 5 min after TSP1 treatment (10 μg/ml) as in **b** in isolated macrophages. Quantification of changes of R123 fluorescence 5 min after oxLDL treatment as in **c**, and 5 min after TSP1 treatment as in **d**. For oxLDL treatment, WT (n=40 for PBS, n=38 for DMSO, n=35 for SSO, n=38 for ACA, n=39 for PJ34, n=44 for U73122 (in HBSS)) and M2KO (n=38 for PBS, n=35 for DMSO) macrophages were from 4 dishes of cultured cells isolated from 3 mice in each group. For TSP1 treatment, WT (n=48 for PBS, n=50 for DMSO, n=53 for SSO, n=51 for ACA, n=57 for PJ34, n=56 for U73122 (in HBSS)) and M2KO (n=48 for PBS, n=52 for DMSO) macrophages were from 4 dishes of cultured cells isolated from 3 mice in each group. (**e-h**) Representative western blot analysis and quantification of the expression of iNOS in isolated macrophages. 3 dishes of cells from 3 mice from each group were chosen for quantification. (**i-l**) Representative real-time Fura-2 Ca^2+^ imaging traces during oxLDL (50 μg/ml) as in **i**, and during TSP1 treatment (10 μg/ml) as in **j**. The averaged traces were from 10 macrophages randomly chosen from a representative culture dish of each group. Quantification of Fura-2 fluorescence changes 5 min after oxLDL treatment as in **k**, and 5 min after TSP1 treatment as in **l**. For oxLDL treatment and TSP1 treatment, 20 macrophages in each group from 3 dishes isolated from 3 mice were chosen for quantification. (**m, n**) Measurement of IL-1β level in culture medium of isolated macrophages after the treatment of oxLDL (50 μg/ml) or TSP1 (10 μg/ml) for 24 h using ELISA. 3 dishes of cells from 3 mice from each group were chosen for quantification. (*: p < 0.05; **: p < 0.01; ***: p < 0.001; ANOVA, Bonferroni’s test; mean ± SEM)

### Inhibiting the activation of TRPM2 in macrophages reduced oxLDL uptake, inhibited macrophage infiltration and improved the impaired macrophage emigration

After confirming that inhibiting TRPM2 activation could suppress macrophage activation, we examined whether these inhibitors affect the phenotypic changes of macrophages induced by oxLDL and TSP1. CD36 inhibitor SSO significantly inhibited the uptake of oxLDL in macrophages derived from bone marrow (Fig. 7a,b), and TRPM2 inhibitor ACA, as well as PJ34 and U73122 also suppressed oxLDL uptake in macrophages (Fig. 7a,b). Moreover, the increased expression of MCP1 and MIF induced by oxLDL or TSP1 was inhibited by ACA, PJ34 and U73122 (Fig. 7c-f), whereas these inhibitors did not produce any suppression of MCP1 and MIF expression in M2KO macrophages subjected to oxLDL (Supplementary Fig. 4a,b) or TSP1 treatment (Supplementary Fig. 4c,d). Furthermore, preincubation with ACA, PJ34 and U73122 inhibited the in vitro macrophage infiltration induced by MCP1 (Fig. 7g,h) and prevented the impairment of emigration ability caused by oxLDL preloading (Fig. 7i,j). These results recapitulate the reduced oxLDL uptake by macrophages, inhibited macrophage infiltration, and preserved macrophage emigration by deleting *Trpm2 in vitro* (Fig 2a-f).

**Figure 7:**
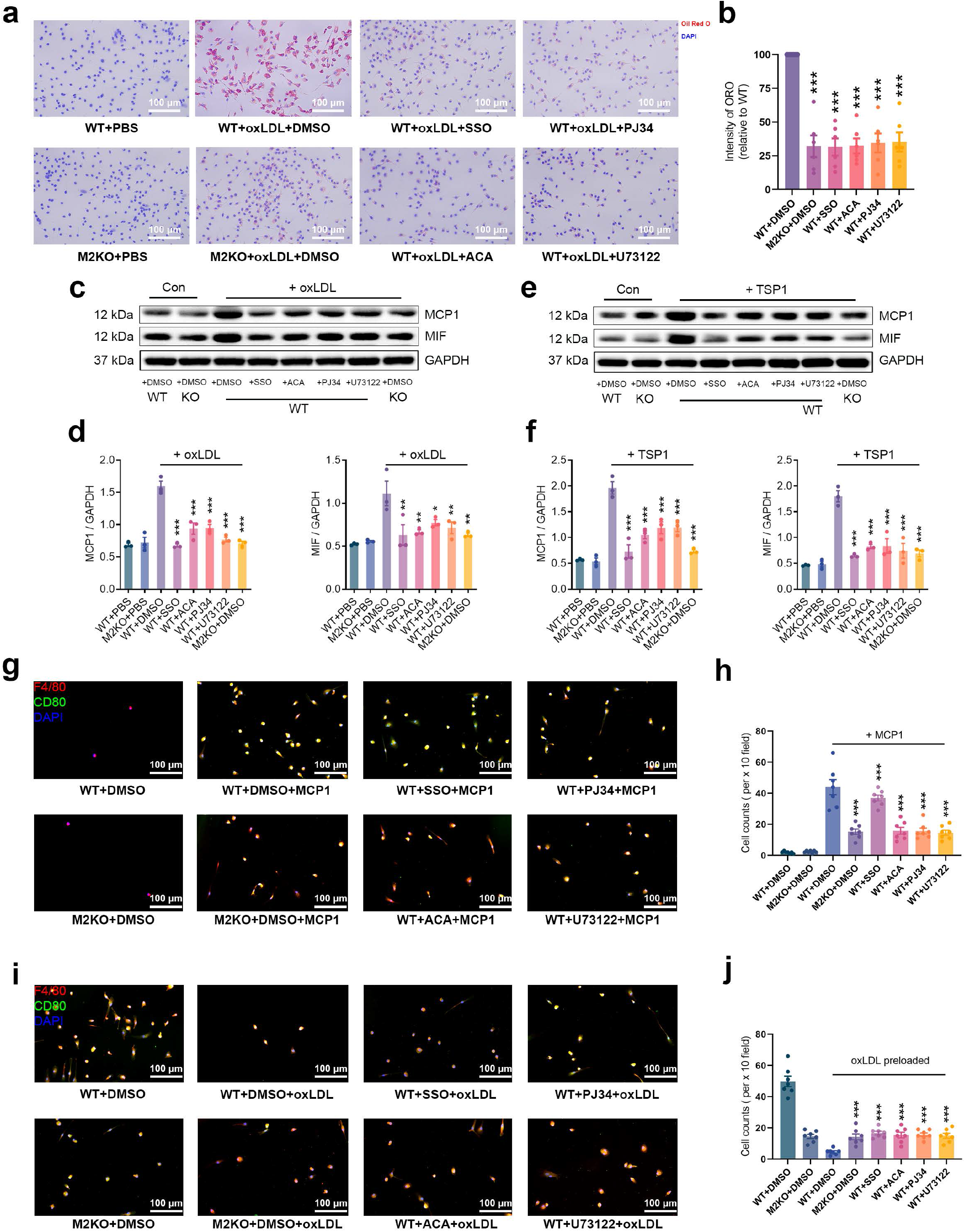
Inhibiting the activation of TRPM2 in macrophages reduces oxLDL uptake, suppresses macrophage infiltration and improves the impaired macrophage emigration caused by oxLDL. (**a, b**) Representative images and quantification of Oil Red O (ORO) staining of isolated macrophages after the treatment with oxLDL (50 μg/ml) for 24 h. 3 dishes of cells from 3 mice from each group were chosen for quantification. (**c-f**) Representative Western blot analysis and quantification of the expression of MCP1 and MIF in isolated macrophages treated with oxLDL (50 μg/ml) or TSP1 (10 μg/ml) for 24 h. 3 dishes of cells from 3 mice from each group were chosen for quantification. (**g, h**) Inhibiting the activation of TRPM2 suppressed macrophage infiltration. *in vitro* macrophage infiltration test was performed as graphic illustration in **Figure 2a** (Red: F4/80; Blue: DAPI; Green: CD80). **h**, Quantification of the number of infiltrated macrophages within a x 10 field. 6 dishes from each group were chosen for quantification. (**i, j**) Inhibiting TRPM2 activation prevented the loss of emigration ability in oxLDL-pre-loaded macrophages. *in vitro* macrophage emigration test was performed as graphic illustration in **Figure 2d**. Macrophage emigration across endothelial cells induced by MCP1. Aorta-derived endothelial cells were plated on the transwell inserts (pore size: 12 μm) for 2-3 days. Bone marrow derived macrophages preloaded with oxLDL for 24 h were added into the upper chamber after endothelial cells completely covered the upper surface of transwells. After 24 h, F4/80 and CD80 staining of macrophages in lower chamber was performed as in **i** (Red: F4/80; Blue: DAPI; Green: CD80). **j**, Quantification of the number of infiltrated macrophages within a x 10 field. 6 dishes from each group were chosen for quantification. (ns: no statistical significance; *: p < 0.05; **: p < 0.01; ***: p < 0.001; ANOVA, Bonferroni’s test; mean ± SEM).

## DISCUSSION

Atherosclerosis is a chronic inflammatory disease with the central pathological features of macrophage infiltration and foam cell formation^1,2^. However, the precise molecular mechanisms regulating these two critical pathogenesis processes remain unclear^2^. Mitigating atherosclerosis is essential for minimizing its complications such as myocardial infarction and ischemic stroke, which are the leading causes of mortality and morbidity^1^. Current available therapies which only control risk factors of atherosclerosis, such as dyslipidermia, have proven effective only to some extents^1^, partially because of the poor patient compliance to lifelong lifestyle modification^41^. Therapeutic strategies which directly target the most important culprit, macrophages, in pathogenesis of atherosclerosis have been lacking due to our incomplete understanding of atherogenic mechanisms. In this study, we revealed that TRPM2 plays a key role in promoting the activation of macrophages in atherogenesis, uncovered the TRPM2-CD36 axis in the pathogenesis of atherosclerosis, and established that targeting TRPM2 in the TRPM2-CD36 axis could be a new therapeutic strategy for atherosclerosis.

TRPM2 is a Ca^2+^-permeable non-selective cation channel activated by oxidative stress^13^, the hallmark of inflammation^42^. With the unique feature of being activated by oxidative stress, TRPM2 has been implicated in several pathological conditions including Alzheimer’s disease, ischemic stroke, inflammatory bowel disease, and inflammatory lung injury^43^. However, whether TRPM2 is involved in atherogenesis was unknown. Here, we demonstrate that *Trpm2* deletion markedly attenuates atherosclerosis in *ApoE*^-/-^ mice fed with HFD. We reveal the mechanism by which *Trpm2* deletion inhibits atherosclerosis is through reducing macrophage burden in atherosclerotic plaque *in vivo*. These findings are supported not only by reduced lesion size and reduced macrophage burden *in vivo*, but also by demonstration of inhibited foam cell formation *in vitro*. Moreover, we discovered an unknown link between TRPM2 and atherosclerosis initiators, the CD36 ligands oxLDL and TSP1. We found that inhibiting TRPM2 markedly suppressed the pro-inflammatory activation of macrophages mediated by oxLDL and TSP1, both of which have been implicated as potent atherogenic activators^37,38,44^. Our discoveries indicate that TRPM2 is a novel therapeutic target for atherosclerosis, and the first direct target of the atherogenesis cascade, in contrast to previous therapies indirectly targeting atherosclerotic risk factors.

The gradual increase of trapped macrophages in the vessel wall during atherosclerosis progression results from two causes: one is the increased number of macrophages infiltrating into the vessel wall, and another is the decreased number of macrophages emigrating back into circulation. By designing two straightforward *in vitro* experiments to mimic *in vivo* conditions, we show that deleting *Trpm2* or inhibiting TRPM2 activation in macrophages not only suppresses macrophage infiltration, but also preserves their emigration ability. MCP1 and MIF, two chemokines produced by activated macrophages themselves, are critical for recruiting macrophages into atherosclerotic plaque^18,20^. We found that the increased expression of MCP1 and MIF induced by oxLDL or TSP1 in macrophages was markedly inhibited by deleting *Trpm2* or inhibiting TRPM2 activation. Uptake of oxLDL significantly inhibits macrophage migration, which was shown to be the major reason for macrophages being trapped inside vessel wall^25^. Our data show that inhibiting TRPM2 activation, or deleting *Trpm2* in macrophages, reduces the uptake of oxLDL, strongly supporting our conclusion that *Trpm2* deletion inhibits foam cell formation.

The uptake of oxLDL is predominately mediated by the scavenger receptor CD36^4^. We found that inhibition of TRPM2 in macrophages *in vitro* or deletion of *Trpm2 in vivo* not only eliminated up-regulation of CD36 as well as CD36-mediated oxLDL uptake, but also largely inhibited downstream signaling cascade of CD36 induced by oxLDL and TSP1, indicating that TRPM2 is necessary for CD36 activation and the subsequent inflammatory responses. Moreover, TRPM2 channel inhibitor ACA, or inhibiting the activation of TRPM2 using PJ34/U73122, exhibited similar effect to that of *Trpm2* deletion, suggesting that TRPM2-mediated Ca^2+^ is necessary for CD36 activation.

One of the intriguing discoveries in this study is the TRPM2 activation mediated by CD36 ligands oxLDL and TSP1, which can be blocked by CD36 inhibitor SSO. The activation of TRPM2 by TSP1 may provide new mechanistic insights about the important role of TSP1 in atherosclerosis shown in a recent study^37^. We uncovered that the mechanisms by which oxLDL and TSP1 activate TRPM2 is through ROS production as measured by R123. ROS production and mitochondrial oxidative stress in macrophages is known to promote macrophage activation in atherosclerotic plaques and accelerate the progression of atherosclerosis^27^. By utilizing R123 real-time cell imaging, we demonstrated for the first time that oxLDL and TSP1 induced a rapid increase of ROS production and mitochondrial oxidative stress in cultured macrophages. The increased ROS and oxidative stress promote ADPR production. As CD36 activation also leads to PLCγ activation which enhances intracellular Ca^2+ 45^, the enhanced ADPR and Ca^2+^ will further activate TRPM2, resulting in a mutually activating feedback loop between TRPM2 and CD36, which perpetuates the inflammatory response in atherosclerosis. To our knowledge, this is the first report demonstrating inter-dependent positive feedback regulating mechanism between TRPM2 and CD36 in promoting atherosclerosis. As CD36 is the most important molecule for oxLDL uptake in macrophages^5^ which results in foam cell formation and inflammatory responses, this new TRPM2-CD36 axis in atherogenesis suggest that TRPM2 may serve as a novel and effective therapeutic target for atherosclerosis.

The TRPM2-CD36 axis constitutes a strong self-promoting mechanism in the initiation and progression of atherosclerosis. We observed a marked TRPM2-mediated intracellular Ca^2+^ increase in macrophages induced by TSP1 or oxLDL. Inhibiting this Ca^2+^ signaling by deleting or inhibiting TRPM2 reduces production of ROS, MCP1 and MIP, inhibits oxLDL uptake, suppresses macrophage infiltration and enhances macrophage emigration. This is the first report demonstrating an important role of TRPM2-mediated Ca^2+^ through TRPM2-CD36 axis in atherosclerosis. Moreover, TRPM2-CD36 mediated Ca^2+^ also promotes NLRP3 inflammasome activation and therefore IL-1β production *in vitro* and *in vivo*, as evidenced that secretion of IL-1β in macrophages induced by oxLDL or TSP1 was markedly inhibited by *Trpm2* deletion, TRPM2 inhibition or blocking the Ca^2+^ release. Consistent with our results, previous studies have implicated that Ca^2+^ is involved in NLRP3 induced inflammasome activation in cultured macrophages^46,47^. Thus, the TRPM2-CD36 activation loop triggers multiple factors and signaling pathways in promoting atherosclerosis. Controlling this atherogenic TRPM2-CD36 axis by targeting TRPM2 provides a promising anti-atherosclerosis strategy.

In conclusion, we found that at the animal level, *Trpm2* deletion protected *ApoE*^-/-^ mice against HFD induced atherosclerosis, which was characterized by reduced plaque burden in the aorta. At the tissue level, *Trpm2* deletion resulted in decreased macrophage burden and suppressed inflammasome activation in the vessel wall. At the cellular level, deletion of *Trpm2* or inhibiting TRPM2 activation in macrophages suppressed macrophage infiltration, decreased oxLDL uptake and improved the impaired macrophage emigration. At the molecular level, oxLDL and TSP1 activated TRPM2 through CD36, and TRPM2 is required for the activation of CD36 signaling cascades in macrophages by oxLDL or TSP1. Moreover, deleting *Trpm2* or inhibiting TRPM2 activation in macrophages inhibited oxLDL-or TSP1-induced ROS production and increase of intracellular Ca^2+^ concentration. Taken together, our studies reveal a novel mechanism for understanding the development and progression of atherosclerosis, and provide a therapeutic strategy that targets on a key player in atherogenesis, TRPM2, for atherogenesis treatment.

## ACKNOWLEDGEMENTS

We thank Dr. Andrew M. Scharenberg (University of Washington) for kindly providing TRPM2 plasmid. This work was partially supported by the National Institute of Health (R01-HL143750) and American Heart Association (19TPA34890022) to LY. CD36-bio-His was a gift from Gavin Wright (Addgene plasmid #52025; http://n2t.net/addgene:52025; RRID:Addgene_52025)^48^.

## AUTHOR CONTRIBUTIONS

L.Y. conceived the research. P.Z. designed and performed in vitro experiments. Z.Y. and J.F. performed most of the *in vivo* experiments. A.S.Y. conducted some in vitro experiments. P.Z. and L.Y. wrote the manuscript with contributions from all the authors.

## METHODS

### Animals Care

All the experimental mice bred and hosted in the animal facility building of University of Connecticut School (UCONN Health) were fed with standard chow diet or high-fat diet (HFD) (Harlan, TD.88137), and water ad libitum. Standard housing conditions were maintained at a controlled temperature with a 12-h light/dark cycle. All experimental procedures and protocols were approved by the Institutional Animal Care and Use Committee (IACUC) of University of Connecticut School of Medicine (animal protocol: AP-200135-0723), and were conducted in accordance with the U.S. National Institutes of Health Guidelines for the Care and Use of Laboratory Animals.

### Knockout of TRPM2 (TRPM2-KO)

The global TRPM2 knockout (*Trpm2*^-/-^) mice were generated by Dr. Yasuo Mori’s lab at Kyoto University, Japan. The deletion of *Trpm2* was developed in C57B6J mice by replacing the third exon (S5–S6 linker in the pore domain) with a neomycin coding region. The knockout mice exhibited no differences in behavior or impairment in breeding, compared to wild type (WT) C57BL/6 mice. *Trpm2*^-/-^ mice were back-crossed to C57BL/6 mice for ≥10 generations before being used for experiments. *Trpm2*^-/-^ mice were crossed with *ApoE*^-/-^ mice (JAX laboratory, 002052) to generate *Trpm2*^-/-^ and *ApoE*^-/-^ mice. Knockout was confirmed by genotyping. The mice were backcrossed with C57BL/6 mice for ≥10 generations before being used for experiments.

### Oil Red O (ORO) staining

Oil Red O (Sigma-Aldrich, O0625) was dissolved in isopropanol at 55 °C for 30 min, and filtered to make a 0.5% stock solution. 30 min prior to use, ORO stock solution was diluted with water at a 6 : 4 ratio, and filtered to make the working solution.

For *in vivo* aorta staining, mice were euthanized based on our animal protocol, and the full-length aorta was carefully dissected out. Aortas were washed 3 times using PBS, and fixed in 10% formaldehyde for 30 min at room temperature. Then aortas were washed 3 times using PBS, and stained by ORO working solution for 5 min at room temperature. After washing 3 times using PBS, the aorta was ready for imaging.

For *in vitro* cultured macrophages staining, mature bone-marrow derived macrophages were plated on 25 mm square coverslips, and treated with oxLDL at a concentration of 50 μg/ml for 24 h. Culture medium was removed and coverslips were washed 3 times using PBS. Macrophages were fixed in 10% formaldehyde for 10 min at room temperature, and washed3 times using PBS. Then macrophages were stained by ORO working solution for 30 s at room temperature, following a wash using 60% isopropanol for 60 s. After washing 3 times using PBS, the coverslip was mounted using Prolong® Gold antifade reagent with DAPI.

### Antibodies, chemicals and reagents

Rabbit polyclonal antibodies to TRPM2 (Novus, NB110-81601, 1:500 in 5% BSA); Rabbit polyclonal antibodies to F4/80 (Santa Cruz Biotechnology, sc-377009-594, 1:100 in 10% goat serum and 5% BSA for immunofluorescence (IF)); Rabbit polyclonal antibodies to CD80 (Santa Cruz Biotechnology, sc-46694-488, 1:1000 in 5% BSA for western blot (WB), 1:100 in 10% goat serum and 5% BSA for IF); Prolong® Gold antifade reagent with DAPI (Life technologies, P36935); Rabbit polyclonal antibodies to MCP-1 (E8Y7P) (Cell Signaling Technology, 81559, 1:1000 in 5% BSA); Rabbit polyclonal antibodies to MIF (E7T1W) (Cell Signaling Technology, 87501, 1:1000 in 5% BSA); Rabbit polyclonal antibodies to CD36 (D8L9T) (Cell Signaling Technology, 14347S, 1:1000 in 5% BSA); Rabbit polyclonal antibodies to Fyn (Cell Signaling Technology, 4023S, 1:2500 in 5% BSA); Rabbit polyclonal antibodies to Phospho-Src Family (Tyr416) (E6G4R) (Cell Signaling Technology, 59548S, 1:2500 in 5% BSA); Rabbit polyclonal antibodies to SAPK/JNK (Cell Signaling Technology, 9252S, 1:2500 in 5% BSA for WB); Rabbit polyclonal antibodies to Phospho-SAPK/JNK (Thr183/Tyr185) (81E11) (Cell Signaling Technology, 4668S, 1:2500 in 5% BSA); Rabbit polyclonal antibodies to p38 MAPK (Cell Signaling Technology, 9212S, 1:2500 in 5% BSA for WB); Rabbit polyclonal antibodies to Phospho-p38 MAPK (Thr180/Tyr182) (Cell Signaling Technology, 9211S, 1:2500 in 5% BSA for WB); Rabbit polyclonal antibodies to iNOS (Santa Cruz Biotechnology, sc-7271, 1:1000 in 5% BSA for WB); Rabbit polyclonal antibodies to GAPDH (Cell Signaling Technology, 7074S, 1:5000 in 5% BSA for WB); HRP-linked anti-rabbit IgG (1:10000 in 5% BSA for WB. NP40 (Thermal Fisher Scientific, 28324), Triton™ X-100 (T-9284), Bovine Serum Albumin (Sigma-Aldrich, 9048-46-8), Goat Serum (Thermal Fisher Scientific, 16210-064). Sulfosuccinimidyl Oleate (sodium salt) (SSO) (Cayman chemical, 11211), N-(p-amylcinnamoyl) Anthranilic Acid (ACA) (Cayman chemical, 14531), PJ-34 (hydrochloride) (Cayman chemical, 14440), U73122 (Cayman chemical, 70740). Recombinant Human Thrombospondin-1 Protein, CF (TSP1) (R&D systems, 3074-TH-050), Recombinant Human CCL2/MCP-1 Protein, (MCP-1) (R&D systems, 279-MC-050/CF). All chemicals for making Tyrode solution and recording solution (see U73122ow) were purchased from Sigma-Aldrich.

### Plasmids and enzymes

CD36 (Addgene, 17928). The pcDNA4/TO-FLAG-hTRPM2 construct was a kind gift from Dr. A.M. Sharenberg(University of Washington, Seattle).

### Cell culture and transfection

HEK293T cells were cultured in Dulbecco’s Modified Eagle’s medium (DMEM) (Thermal Fisher Scientific, 12100-038) supplemented with 10% Bovine Growth Serum (BGS) (HyClone, SH30541.03) and 0.5% penicillin/streptomycin (Thermal Fisher Scientific, 15140-122) at 37 °C and 5% CO2. 8h prior to transfection, culture medium was replaced with DMEM supplemented only with 2.5% BGS. Cells were transfected when at a confluence about 80-90% using Lipofectamine® 3000 Transfection Kit (Thermal Fisher Scientific, 2232162) based on manufacturer’s instruction.

### Isolation and culture of aorta-derived endothelial cell

Endothelial culture medium was made prior to isolation: DMEM: Nutrient Mixture F-12 (DMEM/F12) (Thermal Fisher Scientific, 11330) was supplemented with 100 μg/ml Endothelial cell growth supplement from bovine neural tissue (Sigma, E2759-15MG), 10% Fetal Bovine Serum (FBS) (Thermal Fisher Scientific, A4766) and 0.5% penicillin/streptomycin (Thermal Fisher Scientific, 15140-122).

Wild-type mice were euthanized based on IACUC-approved protocols. Thoracic aorta was quickly dissected out, and lumen was washed 3 times using ice-cold PBS. Then the lumen of the aorta was filled with collagenase II (Worthington, 4177) in DMEM/F12 at a concentration of 1 mg/ml with 2 ends ligated, and digested at 37 °C for 30 min. Then the ligations were released, and homogenate in the aorta was centrifuged at 1000 g for 10 min at 4 °C. The supernatant was carefully removed, and the cell pellet was re-suspended using 20% BSA in DMEM/F12 and centrifuged at 1000 g for 20 min at 4 °C. Then the supernatant was carefully removed, the cell pellet was resuspended with prewarmed endothelial cells culture medium, and cells were plated onto 35 mm culture dishs precoated with Corning® Collagen I, Rat Tail (Corning, 354236). After 24h, medium was replaced and puromycin was added with at a concentration 2 μg/ml (Sigma, P8833-25MG). Culture medium was changed every 2 days. Puromycin was be added in the 1^st^ week to inhibit the growth of other non-endothelial cells. After 1 week, immunofluorescence staining of CD31 was performed to confirm the purity of isolated endothelial cells. Then endothelial cells were plated onto transwell inserts with 12 μm pore size (Costar, 3403) at a density of ~5 × 10^6^ cells/ml. Endothelial cells typically needed 2 ~ 3 days for endothelial cells to completely cover the upper surface of transwell inserts.

### Isolation and culture of bone marrow derived macrophages

Mice were euthanized based on our IACUC-approved protocol, and femurs were quickly removed. Two ends for femurs were cut using a scissor, and bone marrow was washed out using PBS. The collected bone marrow was thoroughly resuspended with DMEM: Nutrient Mixture F-12 (DMEM/F12) (Thermal Fisher Scientific, 11330) supplemented with 25 ng/ml Macrophage Colony-Stimulating Factor from mouse (Sigma, M9170-10UG), 10% BGS (HyClone, SH30541.03) and 0.5% penicillin/streptomycin (Thermal Fisher Scientific, 15140-122). Culture medium was changed every 3 days. After culturing for 7 ~ 9 days, macrophages were usable for experiments. For current recording, Fura-2 and R123 imaging, macrophages were split onto 25 mm square coverslips.

### Isolation of peritoneal macrophages

To elicit macrophage exudation, 1 ml of thioglycolate medium (sodium thioglycolate, 0.5 g/L; yeast extract, 5 g/L; glucose, 5.5 g/L; sodium chloride, 2.5 g/L; L-cystine, 0.5 g/L) was injected intraperitoneally for each mouse. After 4 days, mice were euthanized based on our IACUC-approved protocol. Then abdominal area was sterilized using 70% ethanol, and8 ml of sterilized ice-cold PBS was injected into the peritoneal cavity to collect peritoneal macrophages. Collected peritoneal exudate cells in PBS were centrifuged at 800 g for 10 min at 4°C. Then cells were resuspended in macrophage culture medium and plated on culture dishes at a density of ~2.5 × 10^6^ cells/ml. At 3rd day of isolation, macrophages were subjected to whole-cell TRPM2 current recording.

### Treatment of macrophages

SSO was used to specifically inhibit the activation of CD36 by oxLDL or TSP1. ACA, PJ34 and U73122 were used to inhibit TRPM2 activation by oxLDL or TSP1. SSO, PJ34 and U73122 were dissolved in DMSO at a stock concentration of 10 mM, and ACA was dissolved in DMSO at a stock concentration of 100 mM. They are all diluted to a working concentration of 1 μM in macrophage culture medium or extracellular working solution. U73122 was used in combination with Ca^2+^ free medium and solution. For cell culture, U73122 was diluted in HBSS medium supplemented with BGS and penicillin/streptomycin. For current recording, Fura-2 and Rhodamine-123 imaging, U73122 was diluted in Ca^2+^ free Tyrode solution. For protein extraction, the incubation time for all inhibitors was 8 h. For current recording, and Fura-2 and Rhodamine-123 imaging, inhibitors were added into the extracellular solution to maintain the inhibition.

### *In vitro* macrophage infiltration and emigration test

As shown in the graphic illustration in **Figure 2**, macrophage infiltration and emigration across cultured aorta-derived endothelial cells were examined. Transwell inserts were plated with endothelial cells as described above. For fluid permeation test, ~100000 isolated bone marrow derived macrophages were added into the upper chamber, and recombinant human MCP1 was added into the lower chamber at a concentration of 50 nM to promote macrophage infiltration. For the macrophage infiltration test, macrophages were treated with oxLDL at a concentration of 50 μM for 24h. Then ~100000 oxLDL preloaded macrophages were added into the upper chamber, and recombinant human MCP1 was added into the lower chamber at a concentration of 50 nM to promote macrophage emigration.

### Real-time monitoring of mitochondrial function

Mitochondria function was evaluated using Rhodamine-123 dye quenching as previously reported. Rhodamine-123 (Rh123, Thermal Fisher Scientific, R302) was dissolved in DMSO to make a stock concentration at 10 mg/ml. Pre-warmed DMEM/F12 medium was used to dilute Rhodamine-123 to a 20 μg/ml working concentration. Culture medium was removed and cultured macrophages on the 25 mm coverslip were washed 3 times using prewarmed PBS, then 2 ml of Rh123 working solution was added. Cells were incubated with Rh123 at 37 °C for 15 min. Then Rh123 working solution was replaced with culture medium. Cells were incubated in Tyrode solution for at least 10 min to achieve Rh123 equilibration after the transition of culture medium to Tyrode solution before experiments.

Fluorescence intensities at 509 nm with excitation at 488nm were collected every 15 s for 30 min using CoolSNAP HQ2 (Photometrics) and data were analyzed using NIS-Elements (Nikon).

### Ratio calcium imaging experiments

Changes of intracellular Ca^2+^ were measured using ratio Ca^2+^ imaging as we describe previously. In brief, Fura-2 AM (Thermal Fisher Scientific, F1221) was dissolved in DMSO to make a stock concentration at 1 mM. Pre-warmed DMEM/F12 medium was used to dilute Fura-2 AM to a working concentration at 2.5 μM, and 0.02% Pluronic™ F-127 (Thermal Fisher Scientific, P3000MP) was added to facilitate loading of Fura-2 AM. Macrophages plated on 25 mm glass coverslips were washed 3 times using pre-warmed PBS, and then incubated with 2 ml of Fura-2 AM working solution for 30~45 min at 37 °C. Non-incorporated dye was washed away using HEPES-buffered Saline Solution (HBSS) containing (in mM): 20 HEPES, 10 glucose, 1.2 MgCl_2_, 1.2 KH_2_PO_4_, 4.7 KCl, 140 NaCl and 1.3 Ca^2+^ (pH 7.4).

Ca^2+^ influx was measured by perfusing the cells with Tyrode’s solution under different treatments. Ionomycin (Iono) at 1 μM was applied at the end of the experiment as an internal control. Fluorescence intensities at 510 nm with 340 nm and 380 nm excitation were collected at a rate of 1 Hz using CoolSNAP HQ2 (Photometrics) and data were analyzed using NIS-Elements (Nikon). .

### Western blotting

NP-40/Triton lysis buffer (10% NP40, 1% Triton™ X-100, 150 mM NaCl, 1 mM EDTA, 50 mM Tris, pH=8.0) containing proteinase inhibitors and phosphatase inhibitors was used to lyse both cultured cells and frozen aorta tissue. For cultured macrophages, proteins were extracted 8 h after either oxLDL or treatment with or without different inhibitors. For tissue, full-length aorta were collected 6 months after HFD treatment. Cell and tissue lysate were lysed by ultrasound using an ultrasonic cleaner filled with ice-cold water for 30 min. After incubated on ice for 1 h, lysate was centrifuged at 13000 g for 30 min and supernatant was collected. Protein concentration was measured using Pierce™ Rapid Gold BCA Protein Assay Kit.

50-80 μg of total protein per lane was loaded and separated proteins were transferred to nitrocellulose membranes. Membranes were blocked with 5% BSA and 2.5% goat serum in Tris buffered saline (TBS, pH=7.4) at room temperature for 2 h, and incubated with primary antibodies in TBS with 0.05% Tween (TBS-T) at room temperature for 2 h. Then membranes were incubated with secondary antibodies in TBS-T for 1 h at room temperature before detection. Blots were developed with ImageQuant LAS 4000 imaging system. Band intensity was quantified using ImageJ software and normalized with appropriate loading controls.

### Electrophysiology

Whole cell currents were recorded using an Axopatch 200B amplifier. Data were digitized at 10 or 20 kHz and digitally filtered offline at 1 kHz. Patch electrodes were pulled from borosilicate glass and fire-polished to a resistance of ~3 MΩ when filled with internal solutions. Series resistance (Rs) was compensated up to 90% to reduce series resistance errors to <5 mV. Cells in which Rs was >10 MΩ were discarded^39^.

For heterologous expression, transfected HEK-293 cells were identified by GFP fluorescence. TRPM2 current recording in transfected HEK-293T cells was performed. A fast perfusion system was used to exchange extracellular solutions and to deliver agonists and antagonists to the cells, with a complete solution exchange achieved in about 1–3 s. For recordings using SSO, ACA, PJ34 and U73122, these inhibitors were added into the extracellular recording solution at the same concentration as used during pre-incubation.

Normal Tyrode solution contained (mM): 145 NaCl, 5 KCl, 2 CaCl_2_, 10 HEPES, 10 glucose, osmolarity=290-320 mOsm/Kg, and pH=7.4 was adjusted with NaOH. NMDG-Cl solution contained (mM): 150 NMDG-Cl, 10 HEPES, 10 glucose, osmolarity=290-320 mOsm/Kg, and pH=7.4 was adjusted with NMDG. The internal pipette solution for whole cell current recordings of TRPM2 contained: 135 mM Cs-methanesulfonate (CsSO_3_CH_3_), 8 mM NaCl, 500 nM CaCl_2_, 5 μM EGTA, and 10 mM HEPES, with pH adjusted to 7.2 with CsOH. Free [Ca^2+^]_i_ buffered by EGTA was about 500 nM calculated using Max chelator^39^. ADPR 1 μM was included in the pipette solution for all experiments.

### Immunofluorescence staining

Full length aortas harvested from mice were frozen at −80 °C prior to use, and was mounted in Fisher Healthcare™ Tissue-Plus™ O.C.T. Compound (Thermal Fisher Scientific, 23-730-571) prior to cutting. Aortas were cut into slices at a thickness of 6 μm, mounted to Superfrost® Plus Microscope Slides (Thermal Fisher Scientific, 12-550-15), and frozen at −80 °C for future use. Prior to staining, slides were left at room temperature for at least 30 min to allow for dehydration. Slices were fixed in 10% formaldehyde for 15 min following washing 3 times using PBS, and were incubated in blocking solution containing 5% BSA, 15% goat serum and 1% Triton X-100 at room temperature for 2 h. Primary antibodies were diluted as described previously in TBS-T containing 15% goat serum. Slices were incubated with primary antibodies for at least 12 h at 4 °C following washing 3 times using PBS, and incubated with secondary antibodies at room temperature for 2 h. Then slices were washed 3 times using PBS, mounted using Prolong® Gold antifade reagent with DAPI. Slices were kept at 4 °C before imaging.

### Data analysis

All data are expressed as mean ± SEM. For two groups’ comparison, statistical significance was determined using Student’s t-test. For multiple groups, statistical significance was determined using one-way or two-way analysis of variance (ANOVA), followed by Bonferroni post-test. P<0.05 was considered as significant.

---

**Supplementary Figure 1:**
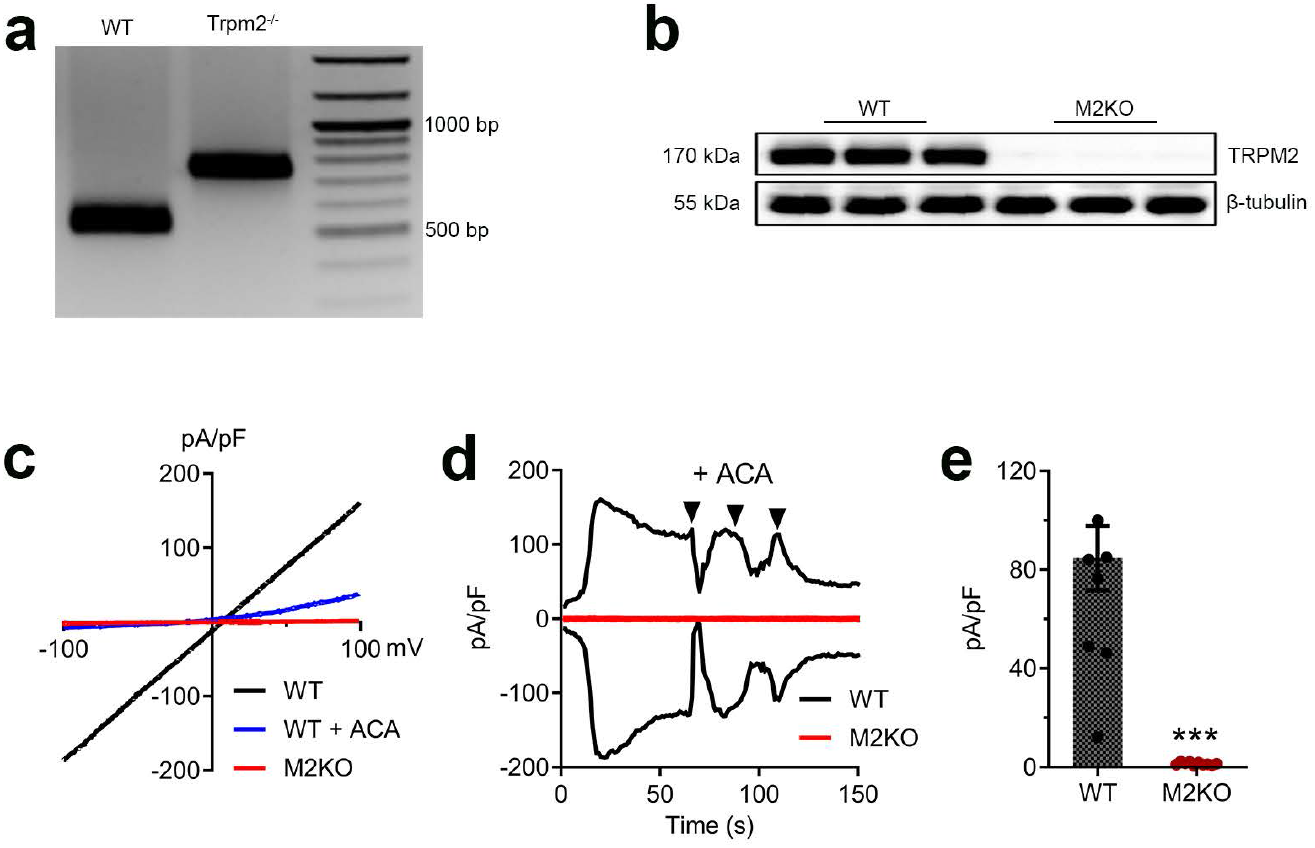
Knockout of *Trpm2* in *ApoE*^-/-^ mice. (**a**) Representative PCR genotyping results showing a 514bp and 740 bp products for WT and M2KO mice. (**b**) Representative Western blot analysis of TRPM2 expression in macrophages isolated from *ApoE* single knockout (WT (n=3)) and *ApoE / Trpm2* double knockout (M2KO (n=3)) mice (**c-e**) Representative recording (**c**, I-V curve; **d**, time-current trace) and quantification of TRPM2 current in macrophages isolated from *ApoE* single knockout (WT) and *ApoE / Trpm2* double knockout (M2KO) mice. ACA is a TRPM2 blocker. (***: p < 0.001; unpaired t test; mean ± SEM)

**Supplementary Figure 2:**
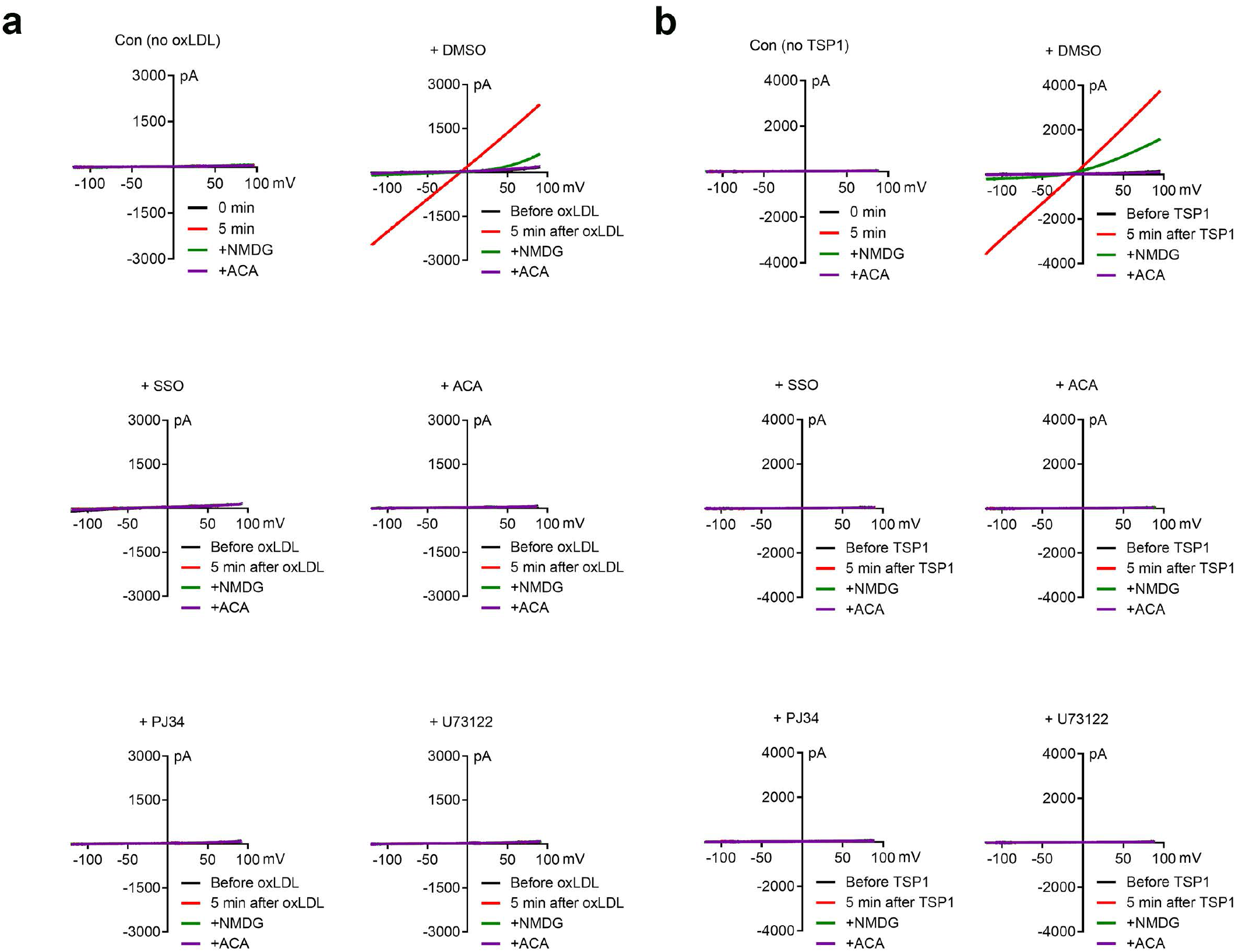
Different inhibitors suppressed the activation of TRPM2 by oxLDL or TSP1 treatment. (**a, b**) Representative recording of TRPM2 current in HEK293T cells transfected with CD36 and TRPM2 during oxLDL treatment (50 μg/ml) as in **a**, and during TSP1 treatment (10 μg/ml) as in **b**. Transfected cells were treated with different inhibitors as indicated before current recording.

**Supplementary Figure 3:**
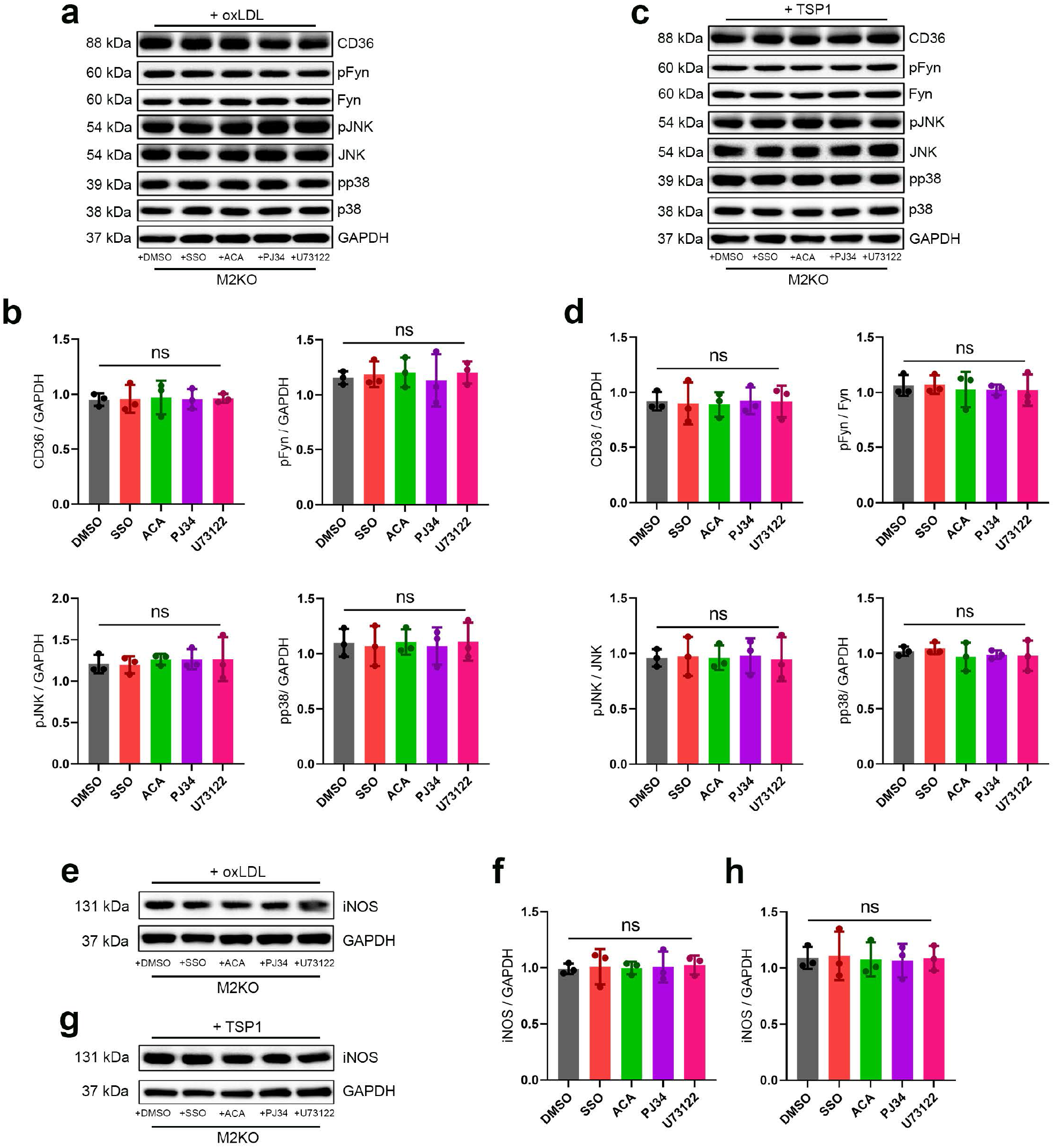
Inhibition of CD36 or TRPM2 did not produce additional inhibitory effect on the activation of CD36 signaling cascade in M2KO macrophages after the treatment of OGD or TSP1. (**a**) Representative Western blot analysis of the expression of CD36, pFyn, Fyn, pJNK, JNK, pp38 and p38 in isolated macrophages from M2KO mice after oxLDL treatment (50 μg/ml). Macrophages were treated with different inhibitors as indicated before protein extraction. (**b**) Quantification of Western blot bands. GAPDH was used for normalization of CD36, Fyn was used for normalization of pFyn, JNK was used for normalization of pJNK, and p38 was used for normalization of pp38. 3 dishes of endothelial cells were used for protein extraction in each group (ns: no statistical significance; mean ± SEM). (**c**) Representative Western blot analysis of the expression of CD36, pFyn, Fyn, pJNK, JNK, pp38 and p38 in isolated macrophages from M2KO mice after TSP1 treatment (10 μg/ml). Macrophages were treated with different inhibitors as indicated before protein extraction. (**d**) Quantification of Western blot bands. GAPDH was used for normalization of CD36, Fyn was used for normalization of pFyn, JNK was used for normalization of pJNK, and p38 was used for normalization of pp38. 3 dishes of endothelial cells were used for protein extraction in each group (ns: no statistical significance; mean ± SEM). (**e**) Representative Western blot analysis of the expression of iNOS in isolated macrophages from M2KO mice after oxLDL treatment (50 μg/ml). Macrophages were treated with different inhibitors as indicated before protein extraction. (**f**) Quantification of Western blot bands. β-tubulin was used for normalization of iNOS (ns: no statistical significance; mean ± SEM). (**e**) Representative Western blot analysis of the expression of iNOS in isolated macrophages from M2KO mice after TSP1 treatment (10 μg/ml). Macrophages were treated with different inhibitors as indicated before protein extraction. (**f**) Quantification of Western blot bands. β-tubulin was used for normalization of iNOS (ns: no statistical significance; mean ± SEM).

**Supplementary Figure 4:**
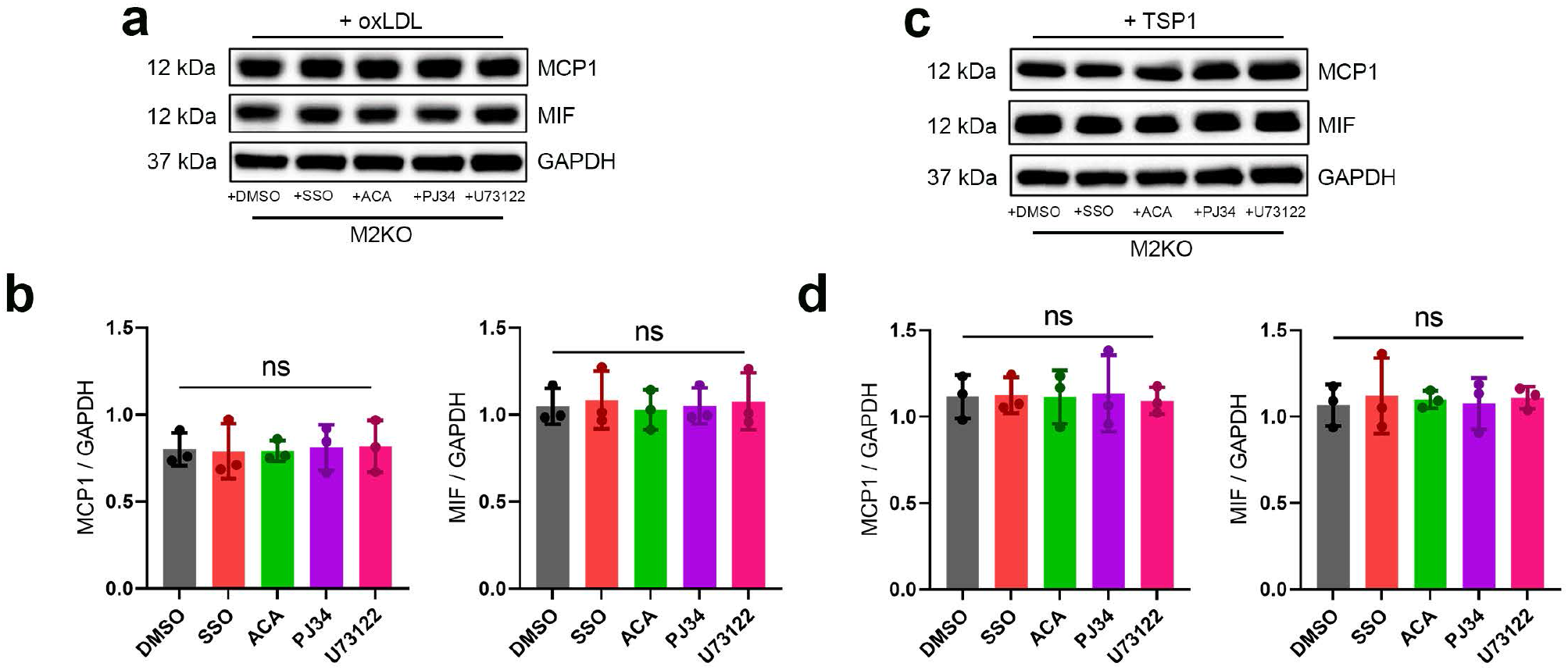
Inhibition of CD36 or TRPM2 did not further inhibit the expression of MCP1 and MIF after the treatment of OGD or TSP1. (**a**) Representative Western blot analysis of the expression of MCP1 and MIF in isolated macrophages from M2KO mice after oxLDL treatment (50 μg/ml). Macrophages were treated with different inhibitors as indicated before protein extraction. (**b**) Quantification of Western blot bands. GAPDH was used for normalization of iNOS (ns: no statistical significance; mean ± SEM). (**c**) Representative Western blot analysis of the expression of MCP1 and MIF in isolated macrophages from M2KO mice after TSP1 treatment (10 μg/ml). Macrophages were treated with different inhibitors as indicated before protein extraction. (**d**) Quantification of Western blot bands. GAPDH was used for normalization of iNOS (ns: no statistical significance; mean ± SEM).

